# Estimating recent and historical effective population size of marine and freshwater sticklebacks

**DOI:** 10.1101/2023.05.22.541730

**Authors:** Xueyun Feng, Ari Löytynoja, Juha Merilä

## Abstract

Effective population size (*N_e_*) is a quantity of central importance in evolutionary biology and population genetics, but often notoriously challenging to estimate. Analyses of *N_e_* are further complicated by the many interpretations of the concept and the alternative approaches to quantify *N_e_* utilising different properties of the data. However, the alternative methods are also informative over different time scales, suggesting that a combination of approaches should allow piecing together the entire continuum of *N_e_*, spanning from the recent to more distant past. To test this in practice, we inferred the *N_e_* continuum for 45 populations of nine-spined sticklebacks (*Pungitius pungitius*) using whole-genome data with both LD- and coalescent-based methods. Our results show that marine populations exhibit the highest *N_e_* values in contemporary, recent, and historical times, followed by coastal and freshwater populations. They also demonstrate the impact of both recent and historical gene flow on *N_e_* estimates and show that simple summary statistics are informative in comprehending the events in the very recent past and aid in more accurate estimation of *N_e_^C^*, the contemporary *N_e_*, as well as in reconstruction and interpretation of recent demographic histories. Although our sample size for large populations is limited, we found that GONE can provide reasonable *N_e_* estimates. However, due to challenges in detecting subtle genetic drift in large populations, these estimates may represent the lower bound of *N_e_*. Finally, we show that combining GONE and CurrentNe2, both sensitive to population structure, with MSMC2 provides a meaningful interpretation of *N_e_* dynamics over time.

## Introduction

By quantifying the magnitude of genetic drift and inbreeding in real-world populations, the concept of effective population size (*N_e_*) has numerous applications in both evolutionary (Charlesworth, 2009; Charlesworth & Charlesworth, 2010) and conservation biology (Frankham et al., 2010; Allendorf et al., 2012). In evolutionary biology, *N_e_* is informative about the efficacy of selection, mutation, and gene flow as systematic evolutionary forces (Charlesworth & Charlesworth, 2010). In conservation biology, it serves as a crucial measure of a population’s evolutionary potential and long-term viability (Frankham et al., 2010; Hare et al., 2011; Waples, 2022a). Various genetic methods have been developed to estimate *N_e_*, ranging from approaches that infer long-term historical *N_e_*dynamics (reviewed in Beichman et al., 2018; Nadachowska-Brzyska et al., 2022) to those designed for estimating contemporary *N_e_* (*N_e_^C^*; Waples & Do, 2010). While significant methodological progress has been made, challenges remain in reconciling different estimators and integrating recent, historical and contemporary *N_e_* estimates into a full *N_e_* continuum, shedding light on how past demographic events shape present-day genetic diversity (Nadachowska-Brzyska et al., 2022). Of the distinct temporal scales, estimating *N_e_^C^*for large populations remains particularly challenging as their genetic drift and inbreeding effects are small and detection of any signal requires extensive sample sizes (Waples et al., 2016; Marandel et al., 2019; Waples, 2024).

In addition to analyses, interpretation of the *N_e_* estimates remains complex (Waples 2024). All methods used to estimate *N_e_* from genetic data make assumptions, and violation of these assumptions may lead to errors and biases (Beichman et al., 2018; Nadachowska-Brzyska et al., 2022, Waples 2024). For instance, many *N_e_* estimation approaches assume populations to be closed and panmictic (e.g., Waples & Do, 2010; but see also Palamara and Pe’er 2013; Santiago et al., 2020; Novo et al., 2023a), while in real life, most populations are structured and affected by at least some level of migration. The methods based on sequential Markovian Coalescent (Li & Durbin, 2011; Schiffels & Durbin, 2014) commonly used to estimate dynamics of the historical *N_e_* are not immune to these effects, and changes in gene flow can yield *N_e_* trajectories that mimic changes in population size (Beichman et al., 2018). Empirical studies examining how introgression and population structure affect *N_e_* estimates across different temporal scales are limited (but see Palamara & Pe’er 2013; Kersten et al., 2023 and Li et al., 2024), highlighting a gap in our understanding of these dynamics.

Marine and freshwater fish populations exhibit distinct genetic variability patterns and demographic histories and offer a compelling case study for investigating the challenges in *N_e_* estimation. Freshwater populations generally harbor less genetic variation than marine populations (e.g., Ward et al., 1994; DeWoody & Avise, 2000; DeFaveri & Merilä, 2015; Kivikoski et al., 2023), indicating smaller long-term *N_e_* (*N_e_^LT^*; Ellegren and Galtier, 2016). This makes intuitive sense due to the limited size and fragmentation of freshwater habitats as compared to more continuous marine environments. However, estimating contemporary *N_e_*for large marine populations is particularly difficult (cf. Waples et al., 2016; Marandel et al., 2019) and rigorous comparisons of marine and freshwater populations are rare (e.g., DeFaveri & Merilä, 2015). Recent development in Linkage Disequilibrium (LD) -based methods (Santiago et al., 2020 & 2024) for the estimation of temporal changes in *N_e_* might bring a solution for this, and the new methods have been shown to provide robust estimates even for populations of relatively large *N_e_* (Santiago et al., 2020 & 2024; Kersten et al., 2023; Atmore et al., 2022 & 2024; Andrews et al., 2024). However, LD-based approaches have biases and limitations of their own (Santiago et al., 2020, Novo et al., 2023a & b, Waples 2024, Gargiulo et al., 2024), particularly their assumption of population isolation that is rarely met in natural populations.

The primary goal of this study was to explore the capability and feasibility of using approaches designed to infer recent and historical changes in *N_e_* to reconstruct the temporal dynamics of *N_e_* of natural populations from different ecological contexts. We focused on nine-spined stickleback (*Pungitius pungitius*), a small euryhaline teleost fish with a circumpolar distribution. The species is present in both open marine and landlocked freshwater habitats and previous studies have found evidence for ancient and potentially ongoing introgression in some parts of its distribution range (Guo et al., 2019; Feng et al., 2022; Wang et al., 2023). Using whole-genome sequences (n = 888) obtained from Feng et al. (2022), we estimated both recent (1–100 generations ago) and historical *N_e_* (hundreds to thousands of generations ago) across 45 marine and freshwater populations originating from localities with varying connectivity and environmental conditions, representing open outbred marine to semi- and fully-isolated inbred freshwater populations.

During the analyses, we observed isolated cases of population structure, prompting us to examine how the violations of panmixia influence the *N_e_* estimates and impact on associated demographic interpretations. More systematically, we leveraged data on admixture between two divergent stickleback lineages (Feng et al. 2022) and investigated how introgression— present at varying levels across populations—influences the temporal dynamics of *N_e_*. By integrating multiple *N_e_* estimators across populations with distinct histories, our study provides complementary insights into reconstruction and interpretation of demographic changes across diverse ecological contexts, as well as the impacts of population structure and gene flow on *N_e_*estimation.

## Methods

### Data Acquisition

The sequence data and admixture proportions used in this study were obtained from a previous study (Feng et al., 2022) and in accordance with the national legislation of the countries concerned. In brief, the data used in this study originate from 45 *P. pungitius* populations covering much of the species distribution area in Eurasia, North America and the Far East (Table S1 and Fig. S1). Of these populations, 12 were from marine and nine from coastal freshwater populations with connection (or recent connection) to the sea. Of the true freshwater populations, eleven were from closed ponds (surface area < 4 ha), ten from lakes, two from rivers and one from a man-made drainage ditch (Table S1). The admixture proportions of nine admixed marine and seven admixed freshwater populations (Table S1) from the Baltic Sea area were derived from Feng et al. (2022).

### Data processing

The short-read data were mapped to the 21 linkage groups (LG) of the v7 nine-spined stickleback reference genome (Kivikoski et al., 2021) using the Burrows-Wheeler Aligner v.0.7.17 (BWA MEM algorithm; Li, 2013) and its default parameters. Duplicate reads were marked with samtools v.1.7 (Li et al., 2009) and variant calling was performed with the Genome Analysis Toolkit (GATK) v.3.6.0 and v.4.0.1.2 (McKenna et al., 2010) following the GATK Best Practices workflows. In more detail, RealignerTargetCreator and IndelRealigner (from v.3.6.0) tools were applied to realign reads around indels, HaplotypeCaller was used to call variants for each individual (parameters set as -stand emit conf 3, -stand call cof 10, - GQB (10,50), variant index type linear and variant index parameter 128000), and finally GenotypeGVCFs was used to jointly call the variants for all the samples using its default parameters. Binary SNPs were extracted with bcftools v.1.7 (Danecek et al., 2021) excluding sites located within identified repetitive sequences (Varadharajan et al., 2019) and negative mappability mask regions combining the identified repeats and unmappable regions (Kivikoski et al., 2021). Sites showing low (<8x) or too high (>25x) average coverage, low (<20) genotype quality, low (<30) quality score and more than 25% missing data were filtered out using vcftools v.0.1.5 (Danecek et al., 2011). Data from the known sex chromosomes (LG12) were removed from further analysis. For details about the subsequent filtering of the dataset used in different analyses, see Table S2.

### Analyses of linkage disequilibrium and genetic relatedness

The magnitude of linkage disequilibrium (LD) and its decay are informative on *N_e_*, level of inbreeding, and migration (Flint-Garcia et al., 2003). Hence, we characterised LD patterns in all populations by estimating the squared correlation coefficient *r^2^* between each pair of SNPs with PopLDdecay (Zhang et al., 2019) with its default settings and max distance between two SNPs were set to 1000Kb. We restricted the analysis to the largest linkage group LG4 and used LG1 for cross-validation (SNP Set 2 in Table S2). The LD decay curve was plotted with R (R Core Team, 2020).

High levels of LD in a population may indicate (i) small *N_e_*, (ii) increased inbreeding, and/or (iii) recent migration/admixture. To distinguish between these, we first estimated the average inbreeding coefficients (*F_IS_*) for each population using vcftools --het. We then used *ngsRelate* v.2 (Hanghøj et al., 2019) to calculate the *r_xy_*, the pairwise relatedness within populations (Hedrick & Lacy, 2015). As a measure of temporal gene flow, we examined the LD decay patterns. Within the same ecotype, populations showing atypical LD decay patterns were considered as potentially affected by temporal gene flow.

### GONE analyses

We reconstructed the recent demographic history using GONE (Santiago et al., 2020). This method utilises the LD patterns in the data and has been shown to be robust for time spans of 0-200 generations before present, even when *N_e_* is relatively large (Santiago et al., 2020). The analyses were performed using the SNP Set1 (Table S2 & S3) along with a genetic map lifted-over from the reference genome version 6 (Varadharajan et al., 2019) to version 7 (Kivikoski et al., 2021). According to Santiago et al. (2020), the possible bias from recent gene flow can be mitigated by lowering the recombination fractions threshold (*hc*). Following that, we repeated the analyses using a *hc* of 0.01 and 0.05. For each population, twenty replicates using the SNP Set1 (Table S2 & S3) were performed with the default settings, and the geometric mean were applied to summarize the *N_e_* estimates across replicates. The replicates are not fully independent due to overlapping SNPs. We used a generation length of two years (DeFaveri et al., 2014) to scale the time and the *N_e_* estimates at 1 generation before present was taken as the estimate of *N_e_^C^* (*N_e_^C^ _Gone_*). Following the patterns shown in Fig. 2F of Santiago et al. (2020), we manually inspected and classified the trajectories as affected by gene flow or not.

### CurrentNe analyses

Although the GONE estimates for the most recent generation can be considered measures of *N_e_^C^*, a newer method called CurrentNe has been shown to provide less biased estimates (Santiago et al., 2024). CurrentNe estimates *N_e_^C^* from the LD patterns between SNPs without requiring information on their location (i.e. genetic map) and accounts for the species’ mating system. The latest version of the method, CurrentNe2 (available at https://github.com/esrud/currentne2), allows for the integration of a genetic map and modelling of gene flow, improving the accuracy of *N_e_^C^* estimation. When the migration option (-x) is enabled, CurrentNe2 assumes the target population to be a metapopulation consisting of two subpopulations of equal size and incorporates migration between them into the *N_e_* estimation. This approach can yield a more appropriate metapopulation estimate in cases where gene flow or population substructure are present (A. Caballero, pers. comm.). We applied CurrentNe2 to estimate the *N_e_^C^ _CurrentNe_* and compared that with *N_e_^C^ _GONE_*. In each population, autosomal SNPs with missing data were removed from the analyses, and the average number of full siblings (k) was empirically inferred from the data. The analysis was performed with and without the migration option, and when population substructure was present, the estimate allowing for migration was taken. The number of SNPs used in this analysis for each population is detailed in Table S4.

### MSMC2 analyses

MSMC2 (Malaspinas et al., 2016) was used to reconstruct the demographic history of the more distant past. As the method can analyse at most eight haplotypes, we selected and utilised the four individuals with the highest sequencing coverage from each population. The input files were generated following Schiffels & Wang (2020), and along with mask files generated by bamCaller.py, the mappability masks (Kivikoski et al., 2021) were applied. Estimates were carried out with default settings, and the outputs were processed assuming mutation rate of 4.37□10^-9^ per site per generation (Zhang et al., 2023) and a generation length of two years (DeFaveri et al., 2014). To conduct bootstrap estimations, the input data were chopped into 1 Mb blocks and an artificial 400 Mb long genome was generated by random sampling with replacement. 20 artificial bootstrap datasets were generated using Multihetsep_bootstrap.py from msmc-tools (https://github.com/stschiff/msmc-tools) and analysed with the same settings as the original data. In all analyses, the first two time segments (which usually are untrustworthy; Schiffels & Durbin, 2014; Sellinger et al., 2021) were discarded. Point estimates of *N_e_* at certain times were obtained by interpolation with R’s *approx* function and the value at 4000 years before present was used as the estimate of *N_e_^H^*, the historical *N_e_*, in statistical analyses.

### Long-term *N_e_* estimation

The average long-term *N_e_* (*N_e_^LT^*) were estimated using the formula (Kimura, 1983) *N_e_*= π*/(*4*µ)*, where *µ* is the mutation rate, assumed to be 4.37□10 per site per generation (Zhang et al., 2023). π was obtained from folded site frequency spectra (SFS), estimated for each population directly from the bam data with ANGSD v.0.921 (Korneliussen et al., 2014), using the R script from Walsh et al., (2022) modified to fit folded SFSs. Sites with more than 70% heterozygote counts were removed and the mappability masks (Kivikoski et al., 2021) were applied in data filtering. For details, see Table S2.

### Statistical analyses

We assessed the linear relationships among alternative *N_e_* estimates and admixture proportions using the Pearson correlation test, implemented in the *cor.test*() function in R (R Core Team, 2020). The Pearson correlation coefficient (*r*) was calculated to quantify the strength and direction of these relationships, with *p*-values reported to assess statistical significance (threshold = 0.05). In cases where normality was violated, we conducted Spearman’s rank correlation as an alternative and the Spearman’s correlation coefficient (*rho*) was reported.

## Results

### Summary statistics show unexpected variation among freshwater populations

We found the genetic diversity to be generally higher in marine and coastal freshwater than in inland freshwater populations and decreasing together with the connectivity of the habitat class (Fig. 1a). However, the patterns of LD decay over physical distance showed marked differences within and between habitat classes (Fig. 1b, Fig. S2). While the decay of LD is faster and shows lower average LD (*r^2^*) in marine than in freshwater populations, the latter show considerable variation, some being similar to marine populations with low levels of LD and others containing very high levels of LD (Fig. 1b). Similarly, we found the marine populations to have generally low *F_IS_*, while those for the freshwater populations were highly variable (Fig. 1c). Since a few populations showed higher than average levels of within-ecotype LD and within-population variation in *F_IS_*, we estimated the pairwise relatedness (*r_xy_*, Hedrick et al. 2015) within each population. The relatedness showed an increase with the degree of isolation of the habitat, and unexpectedly, some freshwater samples were more closely related than the others (*r_x_*_y_ > 0.5 shown in yellow colour in Fig. 1d) within the given site (Fig. S3). The initial analysis revealed that the samples from Lake Riikojärvi, Finland, show an exceptionally high level of LD and strong patterns of inbreeding in comparison to other lake populations, and certain individuals within the population were more closely related than others.

**Fig. 1.**
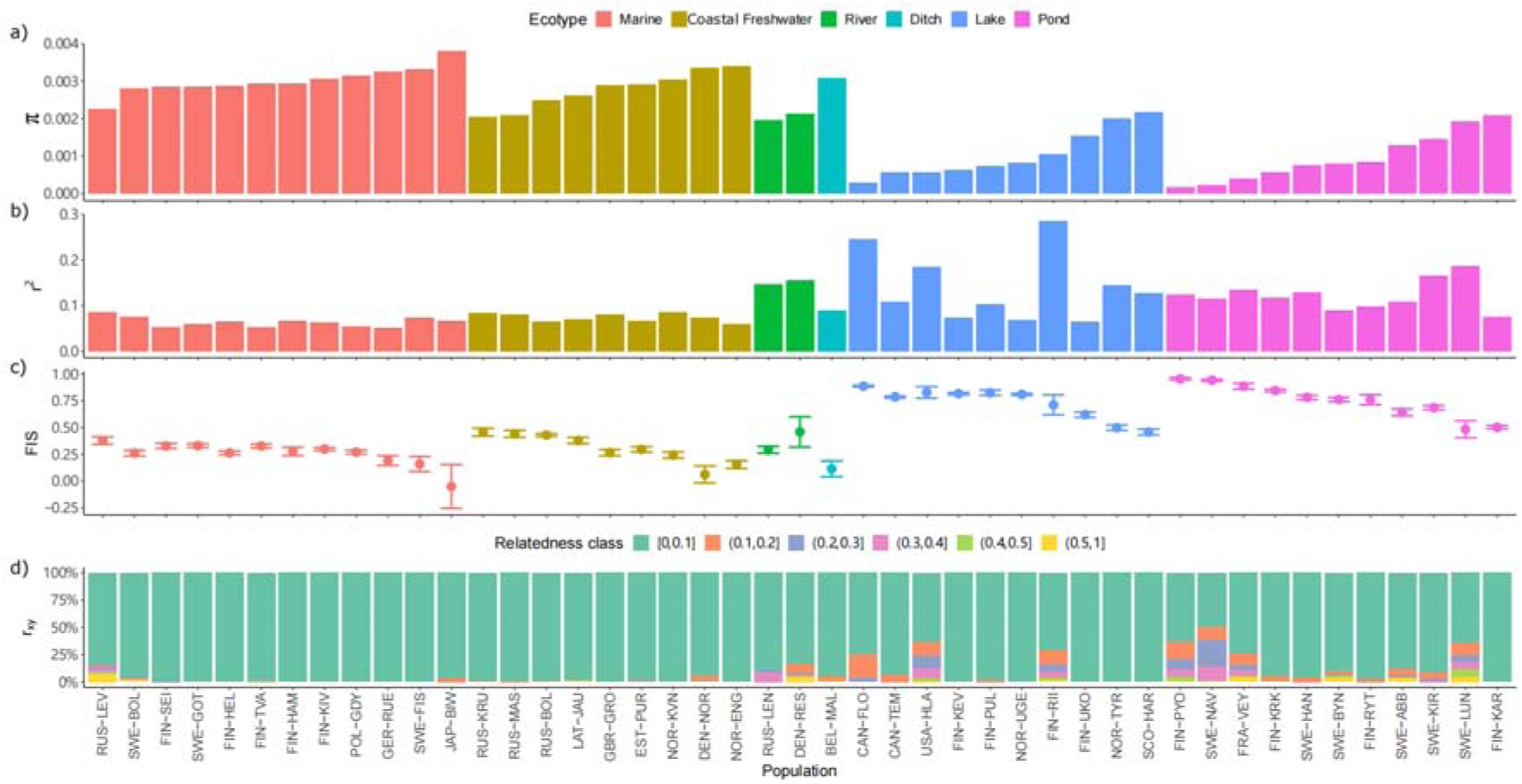
Summary statistics for 45 nine-spined stickleback populations. a) Nucleotide diversity (π) across the autosomal chromosomes. b) LD calculated as the harmonic mean of *r^2^* for SNPs located 100-200 kb apart within LG4. c) Inbreeding coefficients (*F_IS_*) with standard deviations. d) Relatedness (*r_xy_*) for pairs of individuals within populations with colours representing the proportion of pairwise comparisons within a population falling in a specific relatedness class. For the LD decay curve for each population, see Fig. S2.

### Population structure affects inferences of recent demographic history

Population structure, bottlenecks and gene flow can distort the LD patterns and bias the inferences of recent demographic history (Santiago et al., 2020, Novo et al., 2023a). We inferred recent demographic histories using the LD-based methods GONE (Santiago et al., 2020) and found sharp declines in *N_e_* over the last few generations in a few populations (Fig. 2; Fig. S4). Comparisons of *N_e_* estimates across different recombination bin cut-offs (*hc*) showed either inconsistent results or wide ranges of values – the typical symptoms of recent gene flow – for populations exhibiting abnormal LD, inbreeding, or relatedness patterns (Fig. 2; Santiago et al., 2020; Novo et al., 2023a). An extreme case was FIN-RII, an outlier in the LD decay analysis (Fig. 1b, Fig. S2), where *N_e_*estimates ranged dramatically from 14 to 924,485 (Fig. 2) and varied significantly across different *hc* levels (Fig. S4). A lower *hc* value removed the variation in *N_e_* over the generations in a few populations (e.g. FIN-UKO and SWE-GOT), but did not completely eliminate it (e.g. FIN-TVA, Fig. S4). Several populations showed consistent patterns in *N_e_* estimates with different *hc* values. Of the 45 populations, six showed steady *N_e_* estimates over time across the two *hc* values, indicating their stable recent history (Fig. S4).

**Fig. 2.**
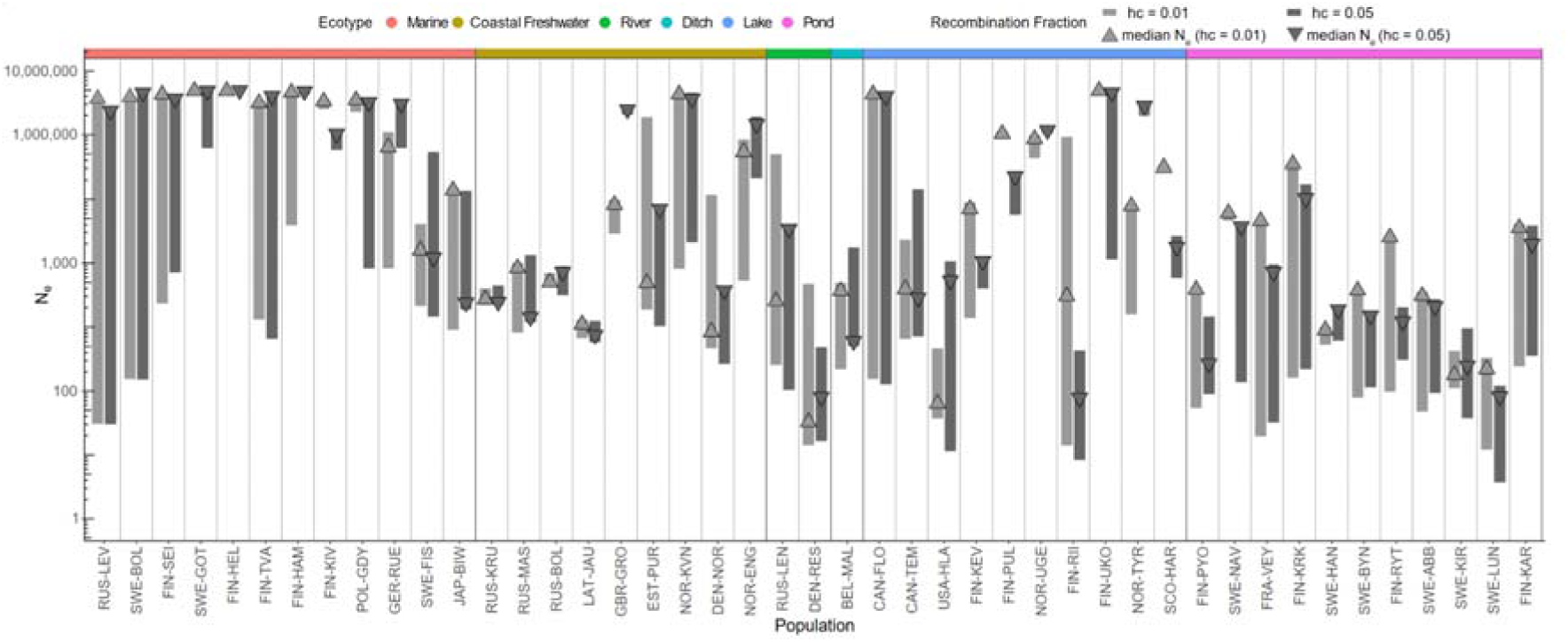
Variations in recent historical *N_e_* estimates obtained with GONE. Range of *N_e_* estimates for each population over 1–50 generations before present, calculated using two recombination fractions (*hc* = 0.01 in light gray, and *hc* = 0.05 in dark gray). Median *N_e_* values are marked by matching triangles. Populations are ordered as in Fig. 1 and the color bar on top represents the ecotypes. For the trajectories of each population, see Fig. S4.

### Differences in *N_e_^C^* estimates

We estimated *N_e_^C^* using the LD-based method CurrentNe2 and compared these with the estimates at 1 generation before present obtained with GONE using different *hc* cut-offs. Overall, the *N_e_^C^* estimates exhibited a similar trend across ecotypes but displayed high variability within freshwater populations, with pond populations generally having smaller *N_e_^C^*s than the others. With CurrentNe2, the *N_e_^C^* of eight marine and seven freshwater populations could not be estimated. Of the 30 remaining populations, 18 were inferred to be affected by migration. As expected, for the 18 populations affected by migration, *N_e_^C^* estimates were higher when migration was accounted for. However, migration rates did not show correlation with the difference (*r* = −0.3, p = 0.23, n = 18) but were weakly correlated (*r* = −0.41, p = 0.09, n = 18) with the ratio of *N_e_^C^* estimates between the two models.

Since we previously showed that the GONE analyses of some populations were influenced by population structure, it is not surprising that the *N_e_^C^_GONE_* estimates for those populations were also affected (10 populations at *hc* = 0.01 and 16 populations at *hc* = 0.05, Fig. 3). *N_e_^C^_GONE_* estimates obtained with *hc* = 0.05 were generally higher than those obtained with *hc* = 0.01, particularly in populations affected by recent gene flow. However, after excluding the affected estimates, the overall trends remained consistent between the two *hc* cut-offs (*r* = 0.50, *p* = 0.006, n = 29). When migration was accounted for in CurrentNe2, we observed a strong alignment between *N_e_^C^ _CurrentNe_* and *N_e_^C^_GONE_* estimates (*hc* = 0.01, *rho* = 0.62, *p* < 0.001, n = 30 and *hc* = 0.05, *rho* = 0.86, *p* < 0.0001, n = 30).

**Fig. 3.**
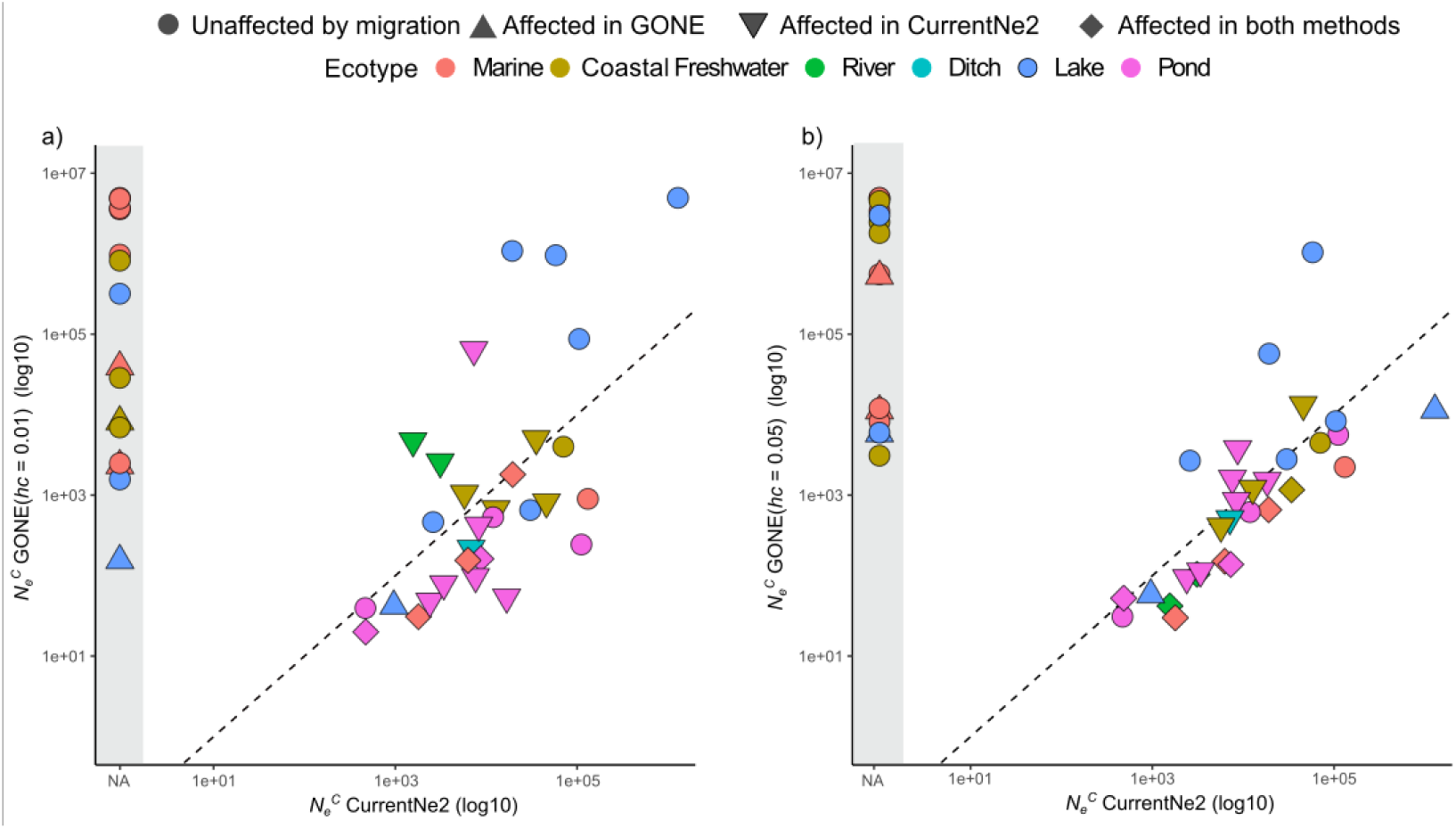
Comparison of estimated contemporary effective population size (*N_e_^C^*) using GONE and CurrentNe2 for 45 nine-spined stickleback populations. a) and b) show *N_e_^C^* estimates obtained from GONE with *hc* = 0.01 and *hc* = 0.05, respectively. The black dashed line represents the 1:1 slope, and the gray shaded area indicates populations for which *N_e_^C^* could not be estimated using CurrentNe2. The shapes denote whether the estimates were affected by migration, while the colors represent the ecotypes.

Discrepancies emerged when comparing *N_e_^C^* estimates between the methods. For example, freshwater populations from Sweden (Lil-Navartjärn, hereafter SWE-NAV) and Finland (Pyöreälampi, hereafter FIN-PYO) that showed high levels of relatedness also obtained markedly different *N_e_^C^* estimates with the two different methods (Fig. S5). A possible explanation is that their exceptionally low genetic diversity and inbreeding have distorted the LD patterns. Substantial differences in *N_e_^C^*estimates were also observed for the Japanese marine population and several Baltic Sea populations (Fig. S5, Table S5). Interestingly, with both methods, the smallest *N_e_^C^*s were observed for a pond population from Lund, Sweden (*N_e_^C^_GONE_* =37.33 at *hc* = 0.01 and 31.30 at *hc* = 0.05, *N_e_^C^_CurrentNe_*=46.63), not in the population with the lowest genetic diversity (FIN-PYO, π = 0.00015), and the highest *N_e_^C^* was observed in a lake population (Ukonjärvi, Finland), followed by a marine population from Biwase Bay, Japan, for CurrentNe2, and the Gulf of Finland for GONE (Fig. S5). In addition, we found that the confidence intervals for *N_e_^C^_GONE_* were extremely narrow compared to those for *N_e_^C^_CurrentNe_*, probably due to the overlapping of SNPs in the replicates (Fig. S5).

### *N_e_* continuums reveal distinctive population histories

We reconstructed the *N_e_* continuum spanning the last 30,000 years, capturing demographic changes from before the Last Glacial Maximum (30 kya) to the present day. Overall, we found both recent historical and historical *N_e_* estimates to be generally lower in freshwater populations compared to marine populations (Fig. 4). However, recent historical *N_e_* estimates exhibit greater variability in lake and coastal freshwater populations compared to pond and marine populations (Fig. 4), consistent with the observed variability in levels of LD and *F_IS_* (Fig. 1), suggesting differences in their recent histories of isolation, recovery from bottleneck or expansion, and migration.

**Fig. 4.**
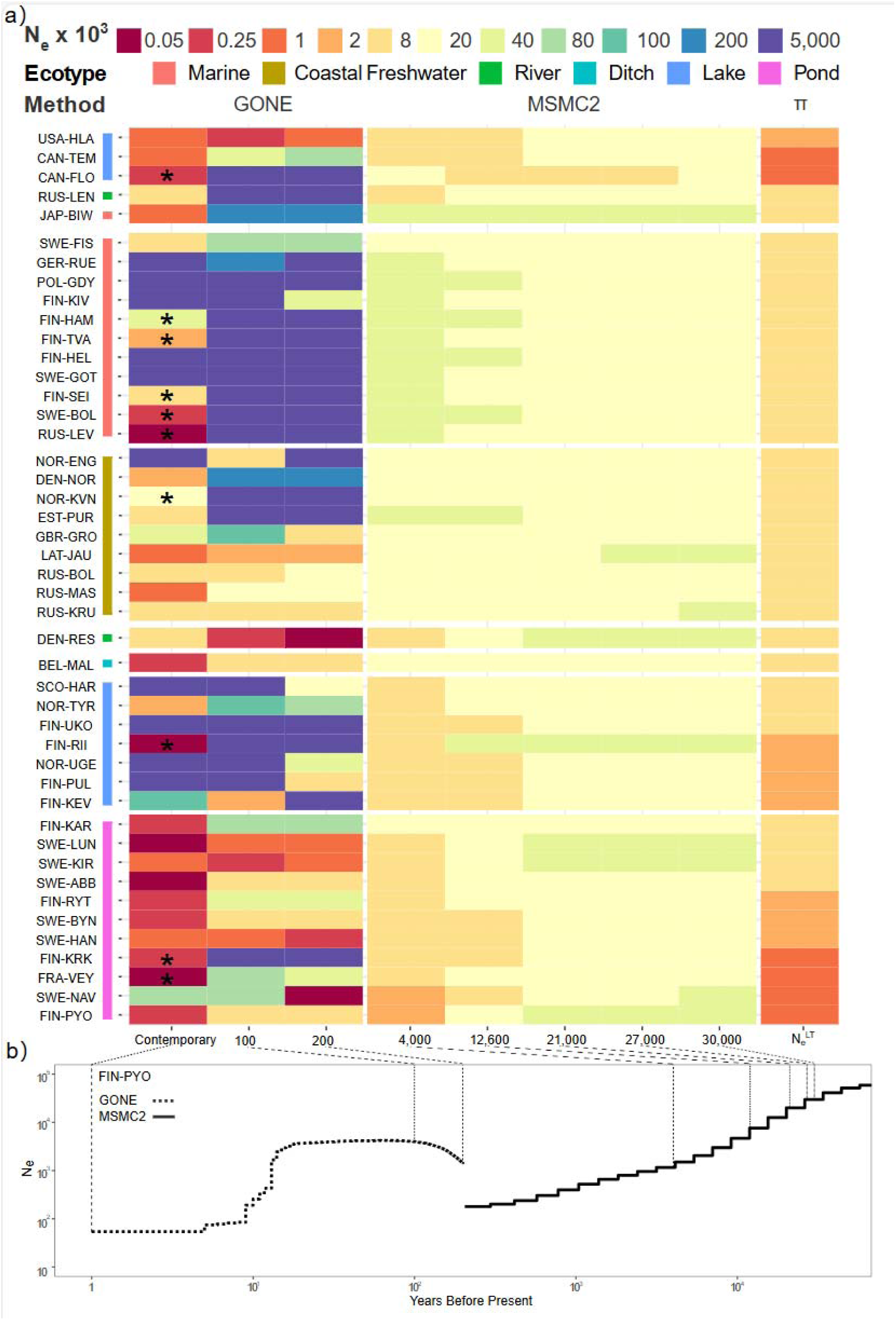
Temporal reconstruction of effective population size (*N_e_*) for 45 nine-spined stickleback populations over the past 30,000 years. a) The colors represent *N_e_* estimates at key time points with the scale given at the top. The estimates for the last 200 years were derived with GONE (*hc* = 0.01; results with *hc* = 0.05 are shown in Fig. S8), while those for 4,000 to 30,000 years ago were obtained with MSMC2. Asterisks indicate estimates influenced by population structure as inferred by manual inspection following the patterns in Fig. 2F of Santiago et al. (2020). Long-term *N_e_* values were calculated from genetic diversity (π). Key time points were chosen to reflect significant geographic events: 4,000 years ago marks the onset of the current Baltic Sea stage, 12,600 years ago the start of the Baltic Ice Lake stage, and 21,000-27,000 years ago corresponds to the Last Glacial Maximum. The color next to population labels represents the ecotypes. b) The demographic history of a representative population, FIN-PYO, over the past 80,000 years. The dashed lines connect the time points in panels a) and b).

The historical *N_e_* show that all populations experienced declines until the end of the Last Glacial Maximum (LGM, approximately 21,000 years ago; Fig. 4, Fig. S6 & S7). After this period, the historical *N_e_* of marine populations increased (Fig. 4, Fig. S7), while most freshwater populations continued to decline (Fig. 4, Fig. S7). Within ecotypes, specific populations exhibit distinct but predictable deviations from the consensus *N_e_* history. For instance, the marine population from Hokkaido, Japan (Fig. S6 & S7), and the freshwater population from Lake Floating Stone, Canada (Fig. S6 & S7), stand out within their ecotypes, reflecting their geographic isolation and the regional climatic differences during the Last Glacial Period. Conversely, some coastal freshwater and pond populations display *N_e_* trajectories similar to those of European marine populations (Fig. S6 & S7), indicating recent colonization of the freshwater environment. In general, the demographic histories of freshwater populations are more variable than those of marine populations (Fig. 4, Fig. S7), reflecting the differing sizes, connectivities, and regional geographic histories of their habitats.

### Historical introgression may differently affect alternative *N_e_* estimators

The Baltic Sea marine populations were inferred to contain 13-30% of Western Lineage ancestry (Feng et al., 2022) and such past introgression events should be visible in the trajectories of historical *N_e_*. The populations show an increase of *N_e_* around 10,000 years ago (Fig. S6 & S7) and similar trajectories are seen in coastal freshwater populations known to contain moderate levels (12-16%) of admixture. However, the Swedish freshwater populations with low levels of admixture (3-5%) in a previous study (Feng et al., 2022) show recent MSMC2 trajectories similar to the consensus pattern of the pond ecotype. This area is known to have been isolated from the Baltic Sea around 10,000 years ago (Mobley et al., 2011), suggesting a negligible impact of low degree of admixture on historical *N_e_* estimates.

To more formally test the impact of historical introgression, we assessed the linear relationships between introgression level and alternative *N_e_* estimators. In the pond populations showing small amounts of admixture, the amount of introgressed ancestry showed no correlation with any *N_e_* estimate (Table 2). Among the Baltic marine populations, the admixture proportion was not correlated with *N_e_^H^*, the *N*_e_ estimate for 4000 years before present, but showed negative correlation (*r* = −0.89, n = 5, p = 0.04) with *N_e_^C^_GONE_* (*hc* = 0.01, Table 2). For the long-term estimates, the effect was opposite and the admixture proportion correlated positively with *N_e_ ^LT^* (Table 2).

**Table 1.** Overview of *N_e_* estimators in this research.

| $N_e$ estimator | Method | Definition |
| --- | --- | --- |
| $N_e^C_{GONE}$ | GONE | Contemporary $N_e$. Estimates at 1 generation before present. |
| $N_e^C_{CurrentNe}$ | CurrentNe2 | Contemporary $N_e$. |
| $N_e^H$ | MSMC2 | Historical $N_e$. Point estimates at 4000 years before present. |
| $N_e^{LT}$ | Genetic diversity | Long-term $N_e$. Estimated from nucleotide diversity ($\pi$) using the equation $N_e^{LT} = \pi / (4\mu)$. |

**Table 2.** Pearson product moment correlations between the different *N_e_* estimates and the admixture proportions in marine and pond ecotypes. Two to nine marine and seven pond populations were included.

| Method | Ecotype | $r$ | $p$ | N |
| --- | --- | --- | --- | --- |
| $N_e^C_{CurrentNe2}$ | Marine | NA | NA | 2 |
| $N_e^C_{GONE}(hc=0.01)$ | Marine | -0.89 | 0.04 | 5* |
| $N_e^C$ GONE ($hc=0.05$) | Marine | -0.12 | 0.85 | 5* |
| $N_e^H$ | Marine | 0.14 | 0.72 | 9 |
| $N_e^{LT}$ | Marine | 0.83 | 0.006 | 9 |
| $N_e^C$ CurrentNe2 | Pond | -0.27 | 0.56 | 7 |
| $N_e^C$ GONE ($hc=0.01$) | Pond | -0.14 | 0.77 | 7 |
| $N_e^C$ GONE ($hc=0.05$) | Pond | -0.28 | 0.55 | 7 |
| $N_e^H$ | Pond | 0.23 | 0.63 | 7 |
| $N_e^{LT}$ | Pond | 0.41 | 0.35 | 7 |

\* Populations that showed drastic decrease in $N_e$ were excluded from analysis.

## Discussion

The aim of this study was to reconstruct the temporal dynamics of *N_e_* in natural populations by employing alternative methods that leverage different genomic signals. Our results revealed that all studied populations had experienced glacial contractions and, while marine populations displayed signs of post-glacial expansion, freshwater populations exhibited continued declines with some populations showing signatures of recovery from historical bottlenecks. Across all estimators, the *N_e_* values were in general highest in marine populations, followed by coastal freshwater populations, and lowest in pond populations. Our analysis also revealed a mixed impact of population structure on *N_e_* estimates. In particular, the *N_e_^LT^* and *N_e_^C^* estimates, based on genetic diversity and obtained with GONE, respectively, for the admixed marine populations correlate with levels of historical genetic introgression, whereas no such correlation was detected in other *N_e_* estimates. We found that characteristic traces in summary statistics can uncover recent gene flow, a violation of assumptions potentially distorting LD patterns and complicating the reconstruction of recent demographic histories, as well as the estimation of *N_e_^C^* with both GONE and CurrentNe2. While also susceptible to historical introgression, MSMC2 models *N_e_* across time and attributes the effects of gene flow to the appropriate historical time periods. We explore these challenges and their implications for accurate and reliable *N_e_* estimation in greater detail below.

### Summary statistics can reveal hidden within-population aberrations

The effective population size *N_e_* has many definitions (Husemann et al., 2016; Waples, 2022) but they all basically return to the Wright-Fisher model (Fisher, 1931; Wright, 1931) and aim to determine the size of an idealised population behaving genetically in a similar fashion to the target population. However, population subdivision and gene flow are common in natural populations (Patton et al., 2019) and such violations of model assumptions impact the inferences of *N_e_* and complicate their interpretation (reviewed in Loog, 2020; Marchi et al., 2021; Waples 2024). In non-model species, limited sample sizes and uncertainty about population conditions pose additional challenges.

Our findings demonstrate that simple summary statistics can be informative in elucidating demographic histories and reduce misinterpretation risks. For example, the Riikojärvi lake population (FIN-RII) was an outlier, exhibiting the highest level of LD, increased relatedness, and highly variable *N_e_* estimates in GONE analyses. These patterns are inconsistent with the population’s nucleotide diversity and the LD levels observed in other populations from the same geographic region and ecotype. We hypothesize that recent hybridization between native fish and genetically distinct migrants may have elevated its LD. Given the isolation of the lake, natural migration seems improbable, implying that these migrants were introduced with fish stocking that has taken place at the lake. Consequently, both *N_e_^C^* estimation and recent demographic reconstructions for this population may be unreliable due to the distortion of LD patterns by recent gene flow.

### Impact of population structure and historical gene flow on N_e_ estimations

The relationship between population’s census size and effective size is complex (Frankham 1995; Palstra & Ruzzante, 2008; Waples et al., 2013). The *N_e_* estimates in this study should be interpreted as indicators of relative demographic changes in each population, and, as such, be indicative of the influence of various demographic processes (e.g., population bottlenecks and inbreeding) and climatic changes. Since the alternative *N_e_* estimators are based on different definitions and utilize distinct features of the data, they probably should not be expected to be fully congruent and none of them should be considered as definitive values of *N_e_*.

Interestingly, we found that gene flow drives divergent responses in *N_e_* estimators: LD-based methods underestimated *N_e_*when migration is not adequately accounted for (Saura et al., 2021; Santiago et al., 2024; Gargiulo et al., 2024), whereas coalescent-based methods tended to overestimate *N_e_*due to the increased genetic diversity introduced by gene flow. This contrast underscores the differences in methodology and sensitivity to demographic changes in different timescales: LD-based methods, which assess *N_e_* based on the variance in reproductive success within recent generations (*N_e_Vk*, Novo et al., 2023a&b, Waples 2024), are sensitive to immediate demographic fluctuations, while coalescent-based methods focus on historical genetic drift and mutation over longer timescales (Hudson, 1990; Li and Durbin, 2011). The distinct temporal scales of the methods emphasize the complexity of interpreting *N_e_*in structured and admixed populations (Kersten et al., 2023) and highlight the importance of integrating multiple *N_e_* estimators to obtain more reliable demographic inferences.

Although GONE is known to be sensitive to population structure (Novo et al., 2023a; Kersten et al., 2023), our results suggest that the method is generally robust across very different-sized populations. The trajectories for marine populations are largely consistent, suggesting that the estimation of recent demographic histories, previously limited to small populations (DeFaveri & Merilä, 2015; Marandel et al., 2019; Nadachowska-Brzyska et al., 2022), may be applied to them as well. However, we also detected deviations and unstable estimates in some populations, particularly those with fluctuating connectivity or showing evidence of population substructure. Such violations of the expected closed population history are known to bias recent historical *N_e_* estimates (1–50 generations before present), often causing sharply decreased *N_e_*estimates (Santiago et al., 2020, Novo et al., 2023a; Kersten et al., 2023; Atmore et al., 2022 & 2024). Limiting recombination between SNPs partially reduced the variation in *N_e_* estimates across generations (Fig. S4; Santiago et al., 2020) but may also lead to a loss of informative SNPs, complicating the reconstruction of recent demographic history (Santiago et al., 2020 & 2023; Novo et al., 2023a) and affecting the estimation of *N_e_^C^*with GONE (Gargiulo et al., 2024).

Consistent with previous observations (Ryman et al., 2019, Novo et al., 2023a&b), our findings indicate that *N_e_^C^*s tend to be underestimated in structured populations. By examining different *hc* cut-offs in GONE, we found that estimates at *hc* = 0.01 were generally lower than those at *hc* = 0.05 but correlated with them, suggesting that reducing the *hc* cut-off might not effectively mitigate the impacts of population structure. On the other hand, lowering the *hc* cut-off reduced the number of informative SNPs and amplified the effects of pseudoreplication, narrowing the confidence intervals (Waples et al., 2022). Furthermore, we found that several populations that GONE inferred unaffected by gene flow showed signals of population structure in CurrentNe2 analyses. The affected populations tended to have lower *N_e_^C^* estimates when migration was not accounted for, with the effect being more pronounced at lower migration rates. Moreover, substantial differences in *N_e_^C^* estimates between the two methods were observed in a few freshwater populations with extremely low genetic diversity. We postulate that the very low genetic variation in these populations causes even minor deviations in the data to appear more pronounced and exaggerate the estimates of changes in drift and LD patterns. Such discrepancies between the methods highlight the importance of careful evaluation of the population structure when estimating *N_e_^C^* and emphasize the need for caution in interpreting the estimates, particularly when the recent conditions of the population are uncertain.

We found that past gene flow had mixed effects on different *N_e_* estimates. In Baltic Sea populations, we observed a positive correlation between *N_e_^LT^* and admixture proportions, suggesting that historical introgression has significantly shaped the genetic landscape of these populations. In contrast, historical *N_e_* estimates sampled at 4000 years before present (ybp), did not show such a correlation. This likely reflects the fact that the ages of the *N_e_^H^*point-estimates are much younger than the recent secondary contacts and the admixture can correctly be taken into account in the more distant time segments. Coalescent-based methods can theoretically accommodate for gene flow and population substructure and are considered relatively robust to low levels of admixture (Beichman et al., 2017; but see Palamara & Pe’er 2013). However, *N_e_* estimates may still be inflated if introgressed genetic variation is not adequately accounted for, as indicated by recent studies on archaic hominins (Li et al., 2024). In the Baltic Sea marine populations, the complex admixture history is mixed with post-glacial population expansion (Feng et al., 2022), and disentangling the effects of these processes on *N_e_* estimates is not straightforward. Utilizing alternative approaches like fastsimcol2 (Excoffier et al., 2021) and dadi (Gutenkunst et al., 2009) might provide a path to a deeper understanding of the history of Baltic Sea populations.

### *N_e_* estimators and demographic history of the nine-spined sticklebacks

While none of the *N_e_* estimates is immune to the impact of population structure and gene flow, they remain valuable for understanding population histories and dynamics. For example, the Japanese marine population from a tide pool connected to Biwase Bay, which experienced inter-species introgression in the distant past (Yamasaki et al., 2020), exhibited the highest *N_e_^LT^*, a median *N_e_^H^*, but a lower *N_e_^C^* compared to Baltic Sea populations. Despite their apparent relative incongruence, the results are consistent with the highly complex history of the population: *N_e_^LT^* reflects the mixed species ancestry with no apparent historical bottleneck and high levels of current genetic diversity, *N_e_*^H^ its marine origin and limited impact during the Last Glacial Period (LGP), *N_e_*^C^ its very recent past in a shallow-water tide pool thereby reduced connectivity to its adjacent marine population and increased genetic drift in the contemporary population. Although anecdotal, the case well represents the differences behind the alternative estimates of *N_e_*, their timescales, differences in sensitivity to demographic changes and associated biological meanings.

The demographic history of nine-spined stickleback populations reflects the significant impact of Pleistocene climatic oscillations. Glaciations and subsequent deglaciations are associated with bottlenecks and expansions, which have shaped the genetic diversity and adaptive potential of contemporary populations (reviewed in Hewitt, 2004). Consistent with other European species (e.g., Backström et al., 2013; Liu et al., 2016), our results show that European nine-spined sticklebacks experienced population contractions during glacial periods. After the LGP (∼11,000 years ago), marine and freshwater ecotypes began to diverge. The timing of bottlenecks in various freshwater populations reflects their colonization history and the formation of new habitats as glacial ice sheets retreated. For instance, pond populations close to the Gulf of Bothnia display bottlenecks 5,000–10,000 years ago, consistent with the historical coastal line in the area (Mobley et al., 2011). In contrast, the White Sea pond populations appear to have resulted from more recent colonizations (Ziuganov & Zotin, 1995), with a bottleneck dated to approximately 500 years ago (Figs. S3 and S4). Additionally, two river populations from the Baltic Sea coast (from Estonia and Latvia) share similar proportions of Western Lineage ancestry with nearby marine populations (Feng et al., 2022), suggesting their establishment occurred after the secondary contact between the two lineages. Such fine details highlight the fundamental differences among the studied freshwater populations, particularly their ages and their origin from ancestral populations representing different pools of standing genetic variation. The latter is a crucial resource for local adaptation (Barrett & Schluter, 2008). Differences in access to the pool of standing variation due to gene flow or historical demography can either constrain or enhance local adaptation (Fang et al., 2021; Kemppainen et al., 2021). In this respect, our results provide a foundation for further investigation into the role of ancestral polymorphism in the local adaptation of nine-spined sticklebacks.

### Feasibility of applying GONE in large populations

In our analyses, estimating the *N_e_^C^* with CurrentNe2 was particularly problematic for certain large populations (e.g. FIN-HEL, n = 22), likely due to the rapid decay of LD and the weak genetic drift signals available for analysis (Gargiulo et al., 2024; Waples, 2025). The latter is inversely proportional to *N_e_*and is quickly diminished by sampling noise as *N_e_* increases (Wang et al., 2016; Waples, 2016). Accurate *N_e_* estimates with LD-based methods require sample sizes comparable to *N_e_* to reduce signal-to-noise ratios to acceptable levels (Hill 1981; Marandel et al., 2019), which is impractical for large populations as this often translates into sampling and sequencing hundreds to thousands of individuals—an unfeasible scale in most cases. In contrast, GONE showed a clear advantage by providing reasonable *N_e_* estimates even with relatively small sample sizes. Furthermore, results from GONE were well aligned with *N_e_^C^*s inferred from CurrentNe2 for both large and small populations when recent gene flow was adequately accounted for. This capability is particularly valuable for estimating *N_e_* of large populations. However, due to the inherent challenges of detecting very subtle drift signals with limited samples, the *N_e_^C^* estimates from GONE might be best viewed to represent the lower bounds of the *N_e_*.

## Conclusions

Using whole-genome data from 45 populations of nine-spined sticklebacks, we demonstrate the utility and limitations of a set of genomic methods for estimating *N_e_* over time. Our study highlights the usefulness of integrating population summary statistics for studying demographic histories and reveals the complex impact of population structure on different *N_e_* estimates. Despite their biases, the estimates remain valuable for gaining a comprehensive understanding of population histories and dynamics. By comparing the results from a method specifically designed to estimate *N_e_^C^*, we demonstrated the potential of using GONE for the estimation of *N_e_^C^* in large populations, which is often a challenge due to their low levels of genetic drift. Overall, our findings underscore the importance of employing a combination of methods to account for both historical and recent demographic processes, providing a more holistic view of population histories and resilience.

## Authors’ contributions

J.M. started the project. X.F., J.M. and A.L. devised the research idea. X.F. performed the analyses with the help of A.L. X.F. and A.L. wrote the first draft and all authors participated in the writing of the final manuscript.

## Acknowledgements

We thank Kirsi Vestenius, Ari Savikko and Kare Koivisto for help with the information of fish transplantation in northern Finland; and numerous collaborators and colleagues for their help in obtaining and processing the samples (listed in Acknowledgements of Feng et al., 2022). The advice and support from Armando Caballero, Andrew Foote, Paolo Momigliano, Petri Kemppainen, and Mikko Kivikoski is gratefully acknowledged. Our research was supported by grants from the Academy Finland (129662, 134728 and 218343 to JM; 322681 to AL), China Scholarship Council (#201608520032 to XF), and Finnish Cultural Foundation (#00210295 to XF). Computational resources provided by the CSC–IT Center for Science, Finland, are acknowledged with gratitude.

## Conflict of interest

The authors declare no competing interests.

## Data availability statement

The whole-genome re-sequencing data have been published previously by Feng et al. (2022) and all the raw sequence data for this study can be accessed through European Nucleotide Archive (ENA) (https://www.ebi.ac.uk/ena) under accession code PRJEB39599. Other relevant data (e.g., input and output files) are available from the Zenodo Open Repository:https://zenodo.org/record/14999855.

## Notes

### Competing Interest Statement

The authors have declared no competing interest.

### Summary of Updates

Manuscript been largely revised to better present the results.

