## Supplemental figs and tables for "Estimating recent and historical effective population size of marine and freshwater sticklebacks": FigS3.pdf

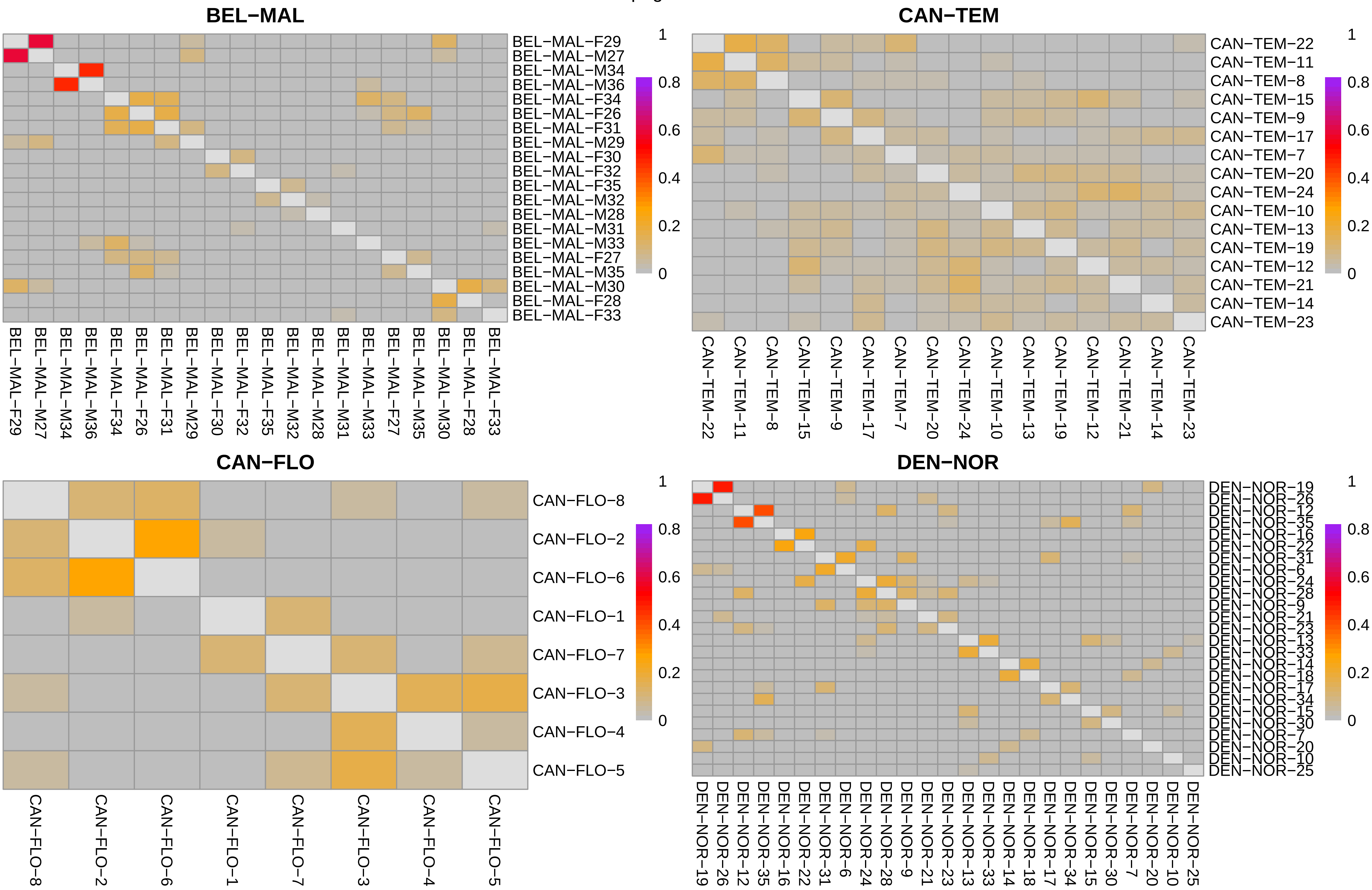
BEL−MAL
BEL−MAL−F29
BEL−MAL−M27
BEL−MAL−M34
BEL−MAL−M36
BEL−MAL−F34
BEL−MAL−F26
BEL−MAL−F31
BEL−MAL−M29
BEL−MAL−F30
BEL−MAL−F32
BEL−MAL−F35
BEL−MAL−M32
BEL−MAL−M28
BEL−MAL−M31
BEL−MAL−M33
BEL−MAL−F27
BEL−MAL−M35
BEL−MAL−M30
BEL−MAL−F28
BEL−MAL−F33
1
0.8
0.6
0.4
0.2
0
CAN−TEM
CAN−TEM−22
CAN−TEM−11
CAN−TEM−8
CAN−TEM−15
CAN−TEM−9
CAN−TEM−17
CAN−TEM−7
CAN−TEM−20
CAN−TEM−24
CAN−TEM−10
CAN−TEM−13
CAN−TEM−19
CAN−TEM−12
CAN−TEM−21
CAN−TEM−14
CAN−TEM−23
1
0.8
0.6
0.4
0.2
0
CAN−FLO
CAN−FLO−8
CAN−FLO−2
CAN−FLO−6
CAN−FLO−1
CAN−FLO−7
CAN−FLO−3
CAN−FLO−4
CAN−FLO−5
1
0.8
0.6
0.4
0.2
0
DEN−NOR
DEN−NOR−19
DEN−NOR−26
DEN−NOR−12
DEN−NOR−35
DEN−NOR−16
DEN−NOR−22
DEN−NOR−31
DEN−NOR−6
DEN−NOR−24
DEN−NOR−28
DEN−NOR−9
DEN−NOR−21
DEN−NOR−23
DEN−NOR−13
DEN−NOR−33
DEN−NOR−14
DEN−NOR−18
DEN−NOR−17
DEN−NOR−34
DEN−NOR−15
DEN−NOR−30
DEN−NOR−7
DEN−NOR−20
DEN−NOR−10
DEN−NOR−25
1
0.8
0.6
0.4
0.2
0

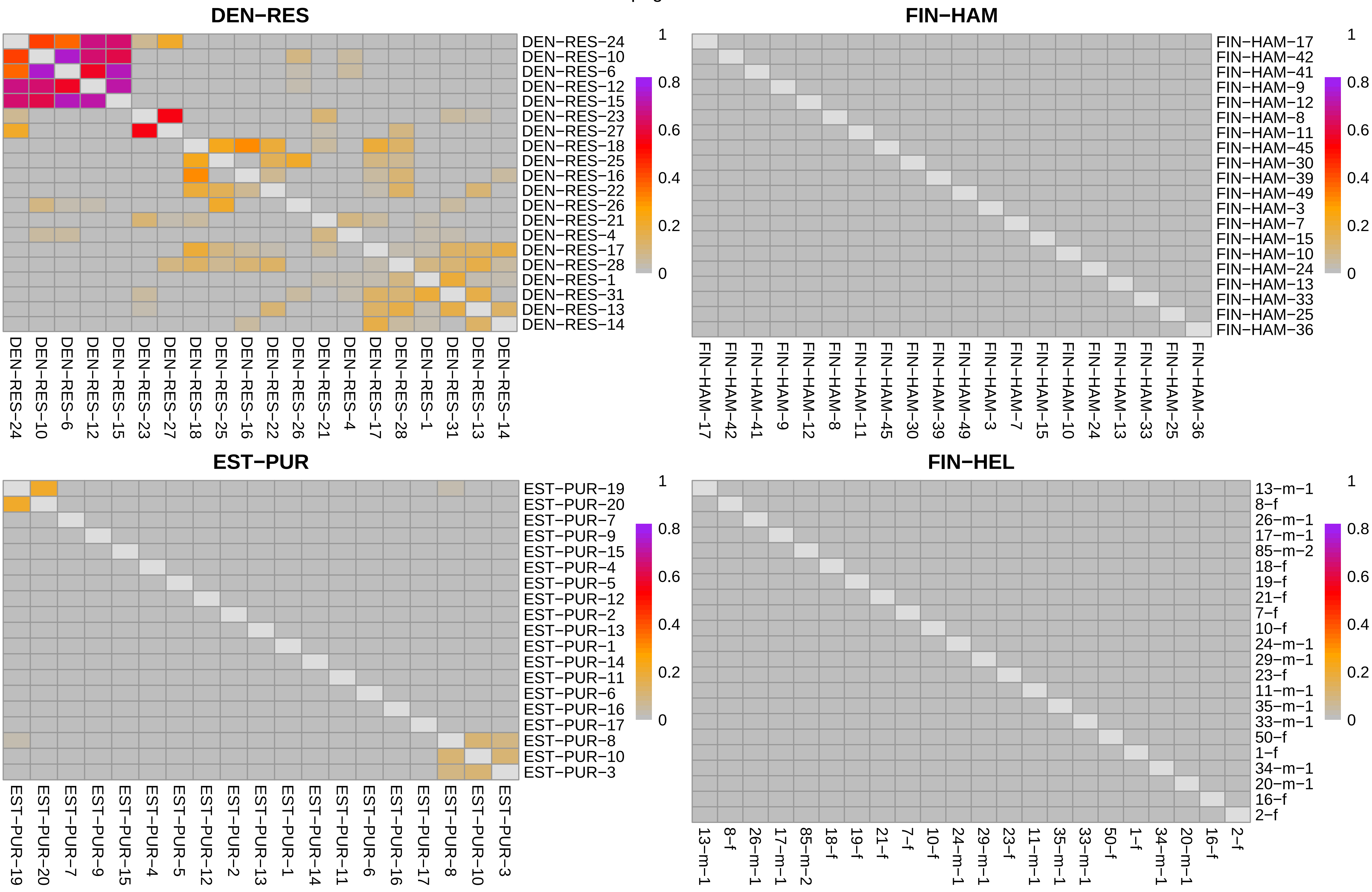
DEN−RES
DEN−RES−24
DEN−RES−10
DEN−RES−6
DEN−RES−12
DEN−RES−15
DEN−RES−23
DEN−RES−27
DEN−RES−18
DEN−RES−25
DEN−RES−16
DEN−RES−22
DEN−RES−26
DEN−RES−21
DEN−RES−4
DEN−RES−17
DEN−RES−28
DEN−RES−1
DEN−RES−31
DEN−RES−13
DEN−RES−14
1
0.8
0.6
0.4
0.2
0
FIN−HAM
FIN−HAM−17
FIN−HAM−42
FIN−HAM−41
FIN−HAM−9
FIN−HAM−12
FIN−HAM−8
FIN−HAM−11
FIN−HAM−45
FIN−HAM−30
FIN−HAM−39
FIN−HAM−49
FIN−HAM−3
FIN−HAM−7
FIN−HAM−15
FIN−HAM−10
FIN−HAM−24
FIN−HAM−13
FIN−HAM−33
FIN−HAM−25
FIN−HAM−36
EST−PUR
EST−PUR−19
EST−PUR−20
EST−PUR−7
EST−PUR−9
EST−PUR−15
EST−PUR−4
EST−PUR−5
EST−PUR−12
EST−PUR−2
EST−PUR−13
EST−PUR−1
EST−PUR−14
EST−PUR−11
EST−PUR−6
EST−PUR−16
EST−PUR−17
EST−PUR−8
EST−PUR−10
EST−PUR−3
FIN−HEL
13−m−1
8−f
26−m−1
17−m−1
85−m−2
18−f
19−f
21−f
7−f
10−f
24−m−1
29−m−1
23−f
11−m−1
35−m−1
33−m−1
50−f
1−f
34−m−1
20−m−1
16−f
2−f

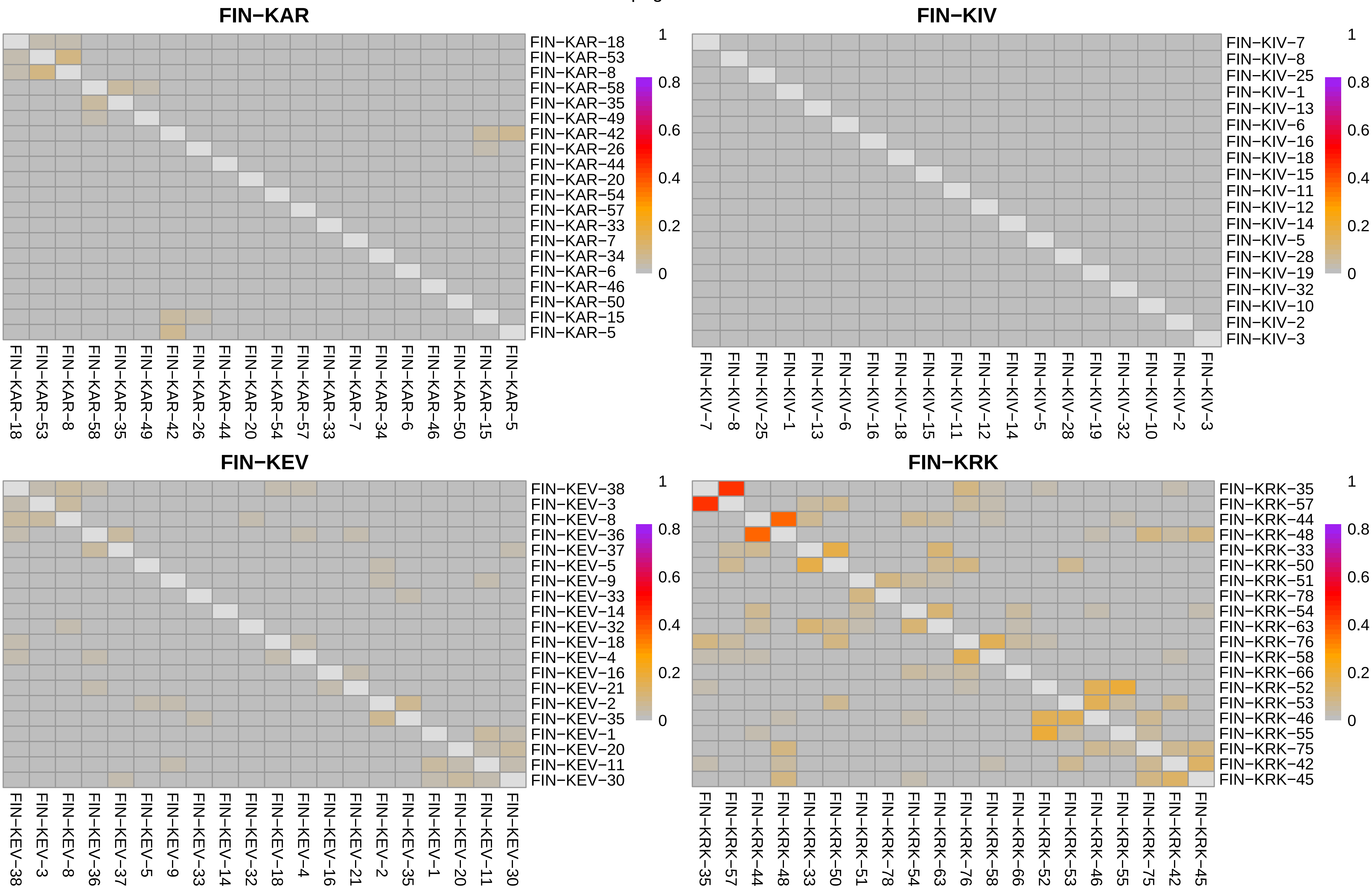
FIN−KAR
FIN−KAR−18
FIN−KAR−53
FIN−KAR−8
FIN−KAR−58
FIN−KAR−35
FIN−KAR−49
FIN−KAR−42
FIN−KAR−26
FIN−KAR−44
FIN−KAR−20
FIN−KAR−54
FIN−KAR−57
FIN−KAR−33
FIN−KAR−7
FIN−KAR−34
FIN−KAR−6
FIN−KAR−46
FIN−KAR−50
FIN−KAR−15
FIN−KAR−5
1
0.8
0.6
0.4
0.2
0
FIN−KIV
FIN−KIV−7
FIN−KIV−8
FIN−KIV−25
FIN−KIV−1
FIN−KIV−13
FIN−KIV−6
FIN−KIV−16
FIN−KIV−18
FIN−KIV−15
FIN−KIV−11
FIN−KIV−12
FIN−KIV−14
FIN−KIV−5
FIN−KIV−28
FIN−KIV−19
FIN−KIV−32
FIN−KIV−10
FIN−KIV−2
FIN−KIV−3
FIN−KEV
FIN−KEV−38
FIN−KEV−3
FIN−KEV−8
FIN−KEV−36
FIN−KEV−37
FIN−KEV−5
FIN−KEV−9
FIN−KEV−33
FIN−KEV−14
FIN−KEV−32
FIN−KEV−18
FIN−KEV−4
FIN−KEV−16
FIN−KEV−21
FIN−KEV−2
FIN−KEV−35
FIN−KEV−1
FIN−KEV−20
FIN−KEV−11
FIN−KEV−30
FIN−KRK
FIN−KRK−35
FIN−KRK−57
FIN−KRK−44
FIN−KRK−48
FIN−KRK−33
FIN−KRK−50
FIN−KRK−51
FIN−KRK−78
FIN−KRK−54
FIN−KRK−63
FIN−KRK−76
FIN−KRK−58
FIN−KRK−66
FIN−KRK−52
FIN−KRK−53
FIN−KRK−46
FIN−KRK−55
FIN−KRK−75
FIN−KRK−42
FIN−KRK−45

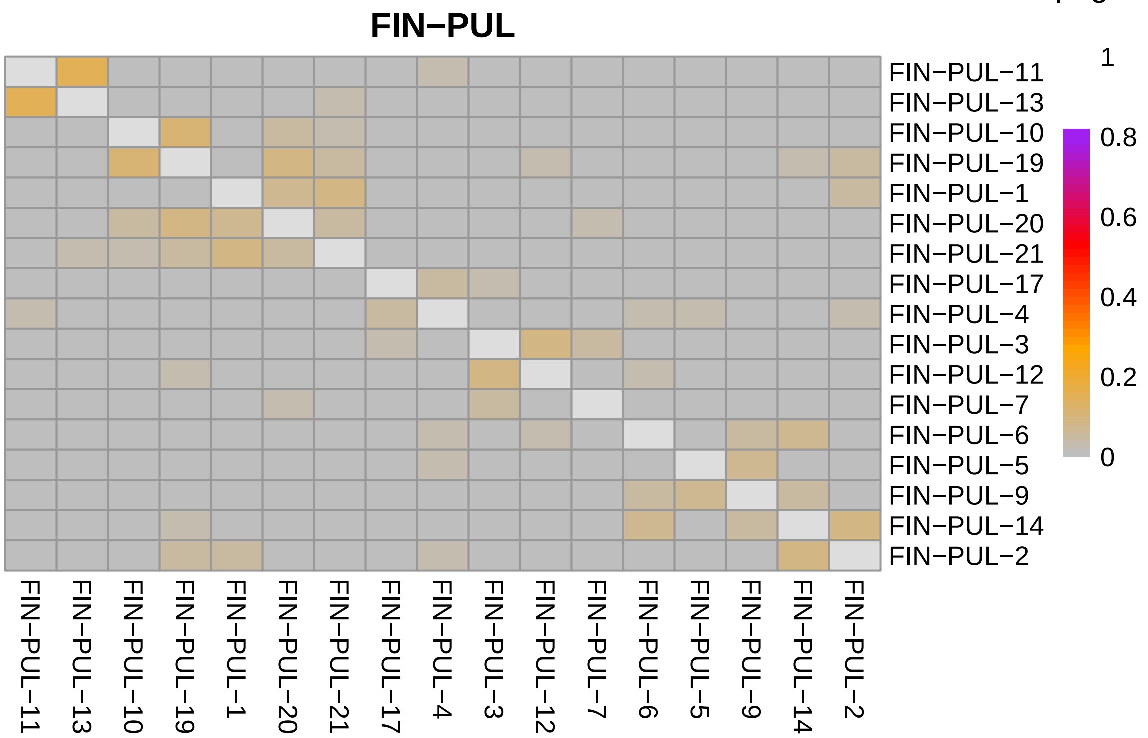
FIN−PUL
FIN−PUL−11
FIN−PUL−13
FIN−PUL−10
FIN−PUL−19
FIN−PUL−1
FIN−PUL−20
FIN−PUL−21
FIN−PUL−17
FIN−PUL−4
FIN−PUL−3
FIN−PUL−12
FIN−PUL−7
FIN−PUL−6
FIN−PUL−5
FIN−PUL−9
FIN−PUL−14
FIN−PUL−2
1
0.8
0.6
0.4
0.2
0

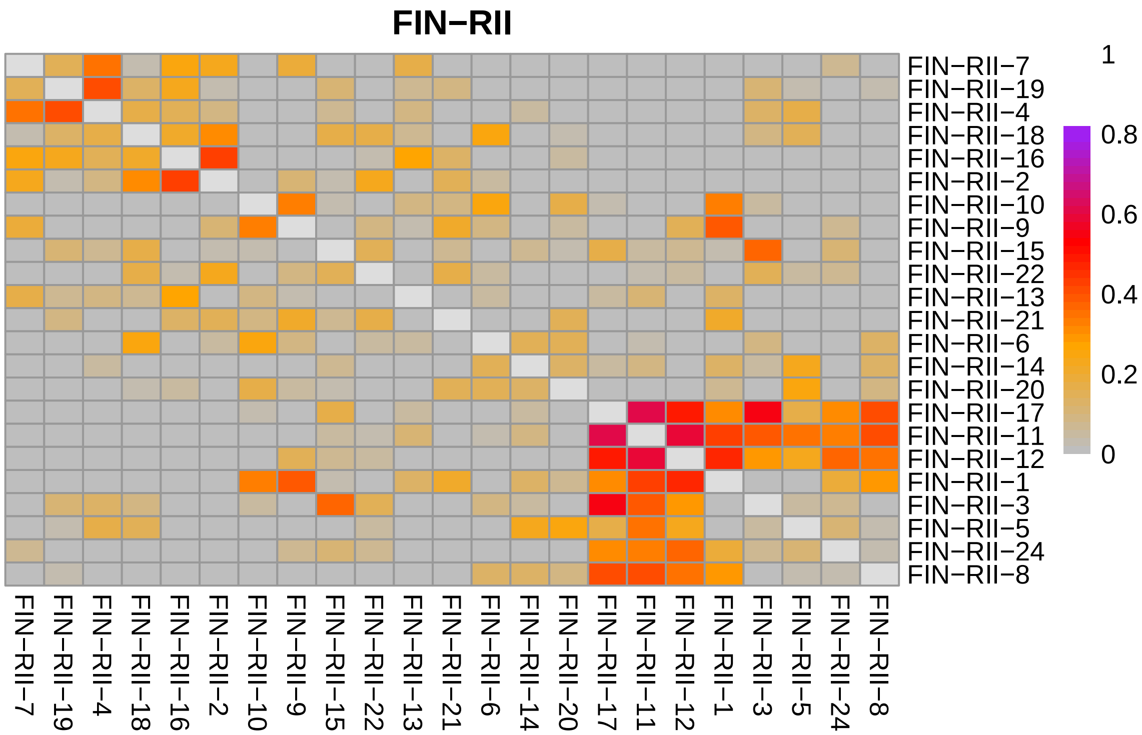
FIN−RII
FIN−RII−7
FIN−RII−19
FIN−RII−4
FIN−RII−18
FIN−RII−16
FIN−RII−2
FIN−RII−10
FIN−RII−9
FIN−RII−15
FIN−RII−22
FIN−RII−13
FIN−RII−21
FIN−RII−6
FIN−RII−14
FIN−RII−20
FIN−RII−17
FIN−RII−11
FIN−RII−12
FIN−RII−1
FIN−RII−3
FIN−RII−5
FIN−RII−24
FIN−RII−8
1
0.8
0.6
0.4
0.2
0

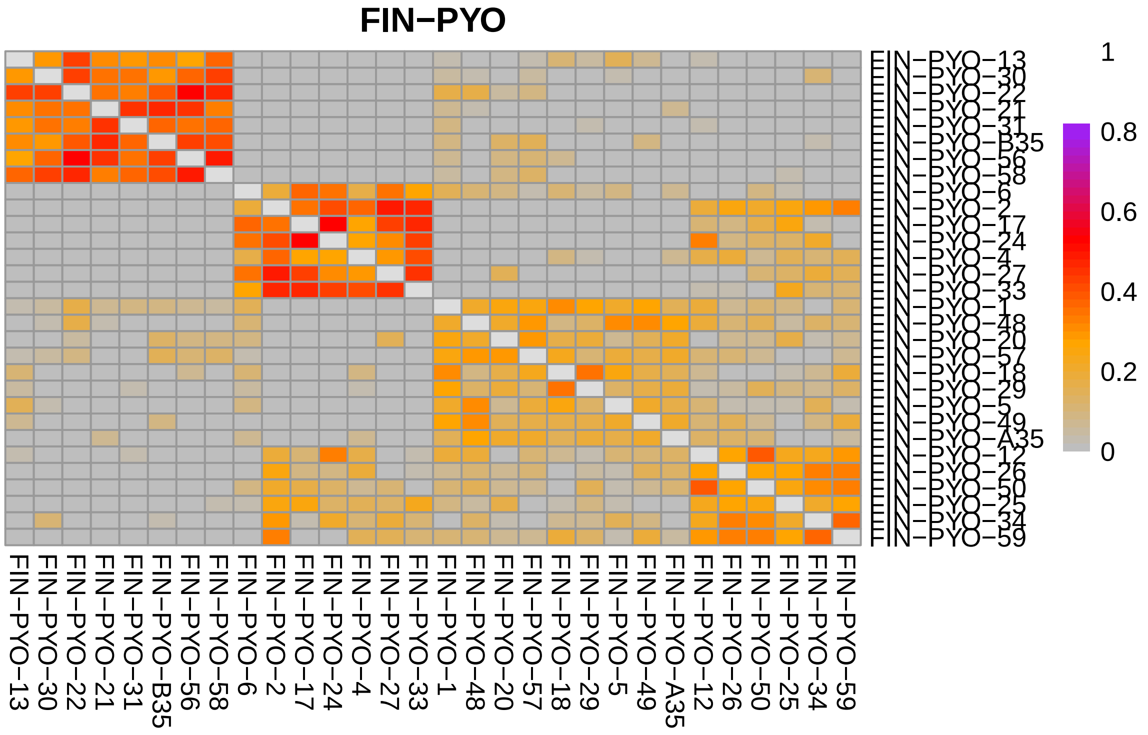
FIN−PYO
FIN−PYO−13
FIN−PYO−30
FIN−PYO−22
FIN−PYO−21
FIN−PYO−31
FIN−PYO−B35
FIN−PYO−56
FIN−PYO−58
FIN−PYO−6
FIN−PYO−2
FIN−PYO−17
FIN−PYO−24
FIN−PYO−4
FIN−PYO−27
FIN−PYO−33
FIN−PYO−1
FIN−PYO−48
FIN−PYO−20
FIN−PYO−57
FIN−PYO−18
FIN−PYO−29
FIN−PYO−5
FIN−PYO−49
FIN−PYO−A35
FIN−PYO−12
FIN−PYO−26
FIN−PYO−50
FIN−PYO−25
FIN−PYO−34
FIN−PYO−59
1
0.8
0.6
0.4
0.2
0

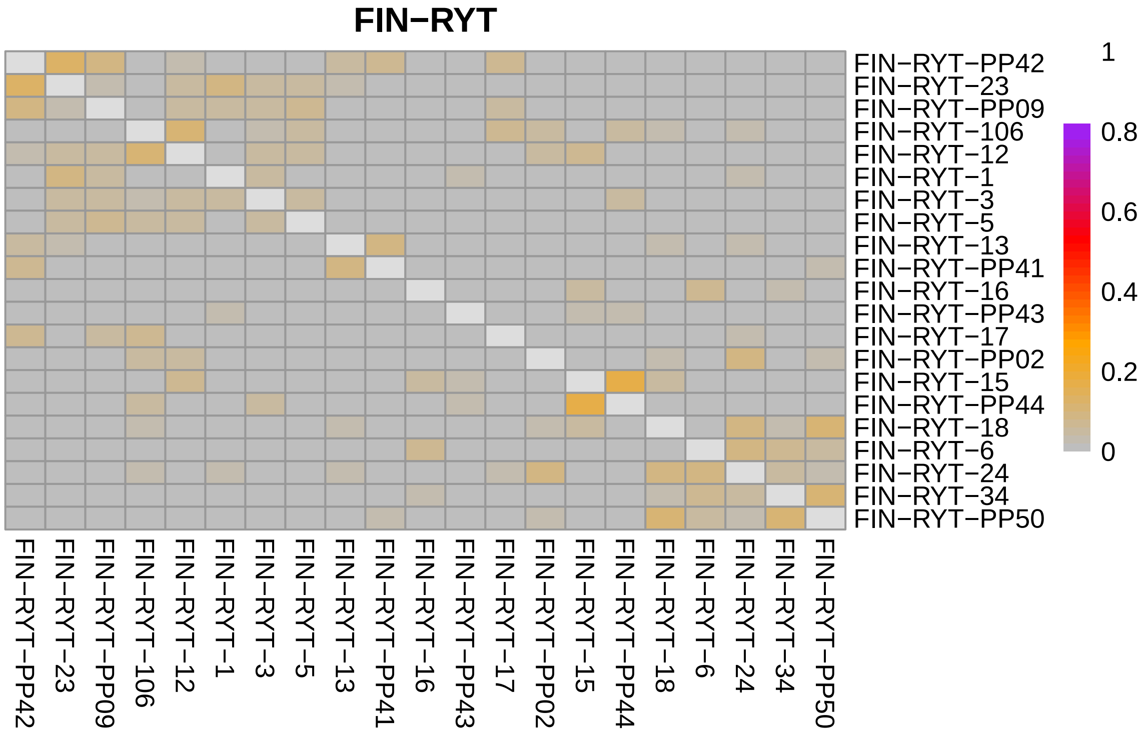
FIN−RYT
FIN−RYT−PP42
FIN−RYT−23
FIN−RYT−PP09
FIN−RYT−106
FIN−RYT−12
FIN−RYT−1
FIN−RYT−3
FIN−RYT−5
FIN−RYT−13
FIN−RYT−PP41
FIN−RYT−16
FIN−RYT−PP43
FIN−RYT−17
FIN−RYT−PP02
FIN−RYT−15
FIN−RYT−PP44
FIN−RYT−18
FIN−RYT−6
FIN−RYT−24
FIN−RYT−34
FIN−RYT−PP50
1
0.8
0.6
0.4
0.2
0

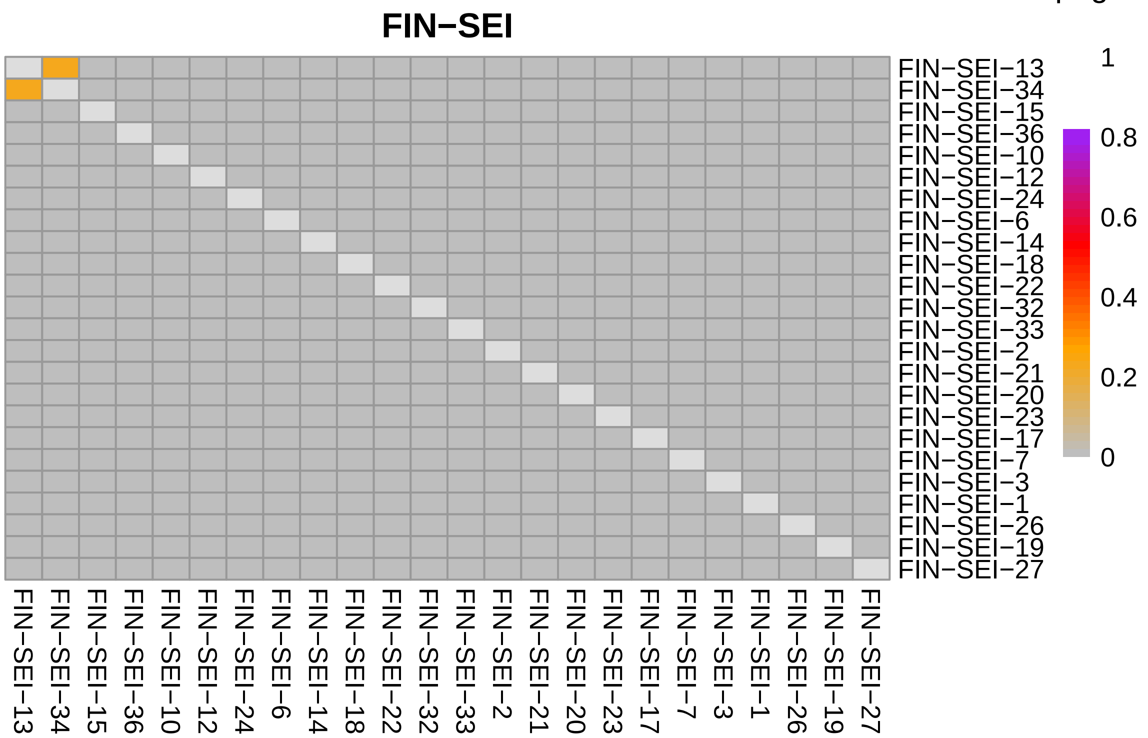
FIN−SEI
1
0.8
0.6
0.4
0.2
0
FIN−SEI−13
FIN−SEI−34
FIN−SEI−15
FIN−SEI−36
FIN−SEI−10
FIN−SEI−12
FIN−SEI−24
FIN−SEI−6
FIN−SEI−14
FIN−SEI−18
FIN−SEI−22
FIN−SEI−32
FIN−SEI−33
FIN−SEI−2
FIN−SEI−21
FIN−SEI−20
FIN−SEI−23
FIN−SEI−17
FIN−SEI−7
FIN−SEI−3
FIN−SEI−1
FIN−SEI−26
FIN−SEI−19
FIN−SEI−27

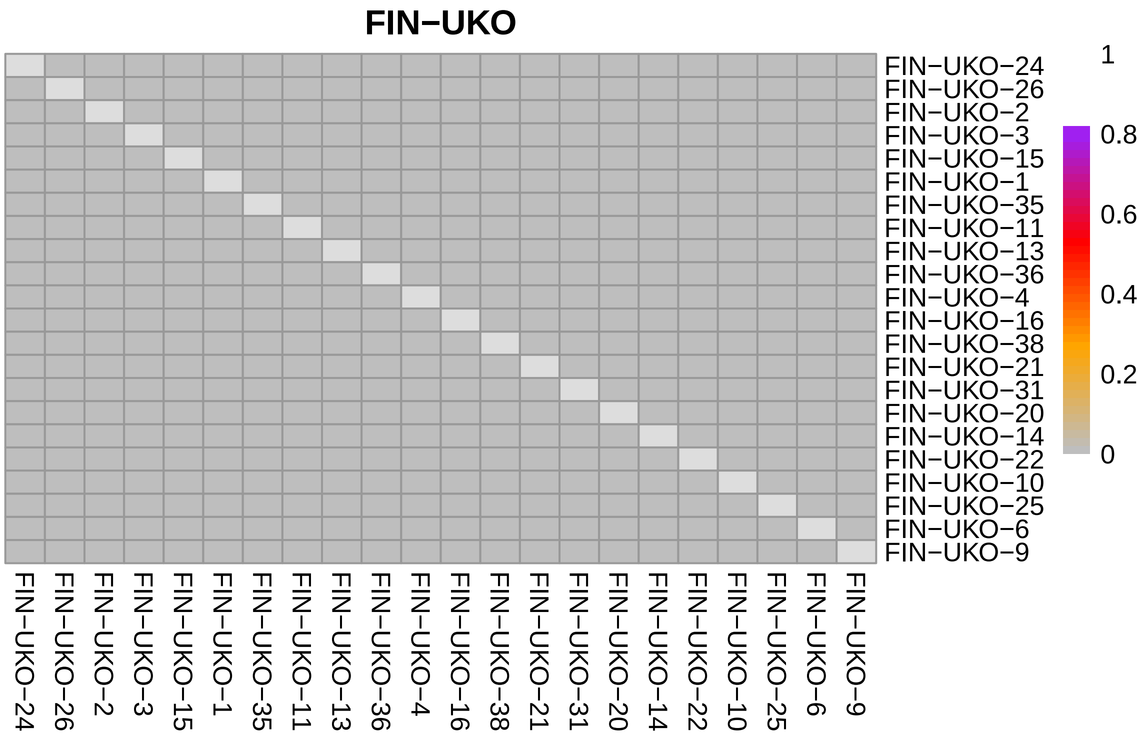
FIN−UKO
1
0.8
0.6
0.4
0.2
0
FIN−UKO−24
FIN−UKO−26
FIN−UKO−2
FIN−UKO−3
FIN−UKO−15
FIN−UKO−1
FIN−UKO−35
FIN−UKO−11
FIN−UKO−13
FIN−UKO−36
FIN−UKO−4
FIN−UKO−16
FIN−UKO−38
FIN−UKO−21
FIN−UKO−31
FIN−UKO−20
FIN−UKO−14
FIN−UKO−22
FIN−UKO−10
FIN−UKO−25
FIN−UKO−6
FIN−UKO−9

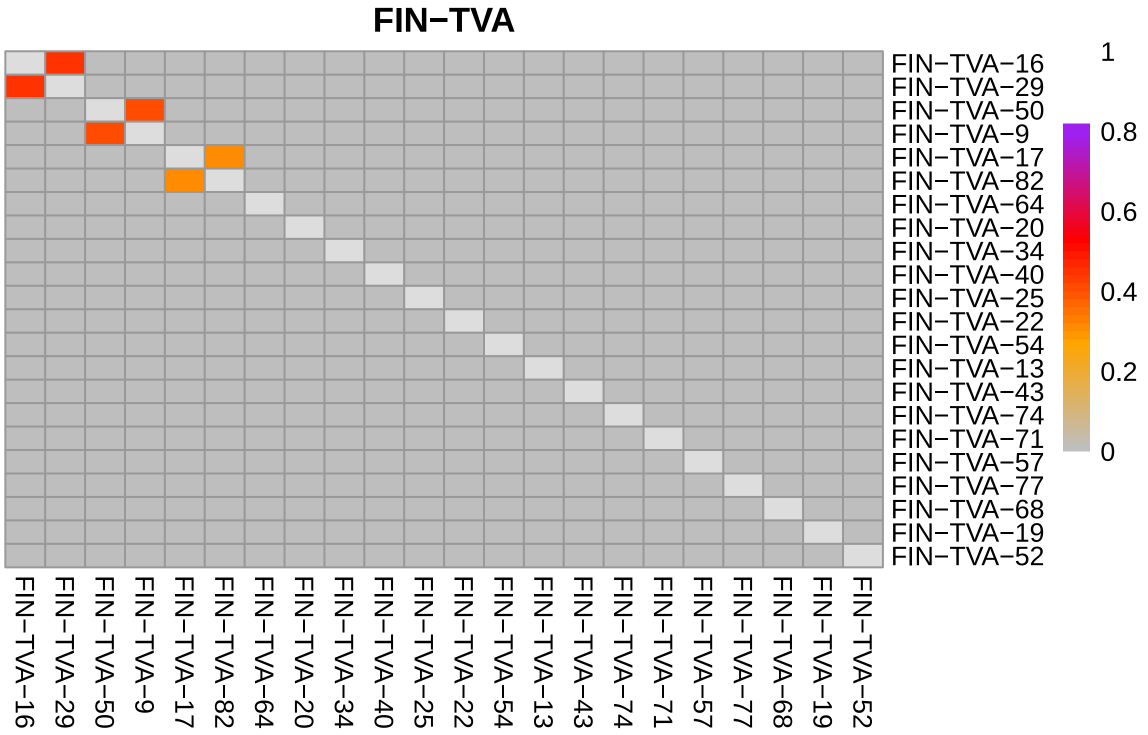
FIN−TVA
1
0.8
0.6
0.4
0.2
0
FIN−TVA−16
FIN−TVA−29
FIN−TVA−50
FIN−TVA−9
FIN−TVA−17
FIN−TVA−82
FIN−TVA−64
FIN−TVA−20
FIN−TVA−34
FIN−TVA−40
FIN−TVA−25
FIN−TVA−22
FIN−TVA−54
FIN−TVA−13
FIN−TVA−43
FIN−TVA−74
FIN−TVA−71
FIN−TVA−57
FIN−TVA−77
FIN−TVA−68
FIN−TVA−19
FIN−TVA−52

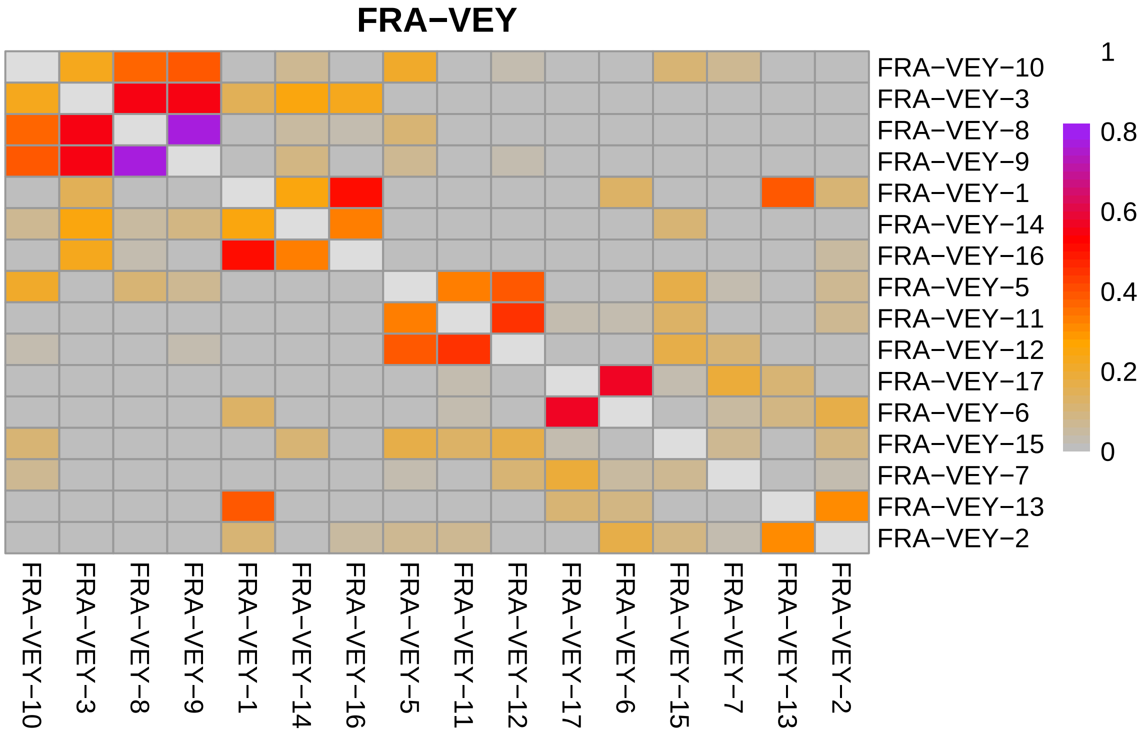
FRA−VEY
1
0.8
0.6
0.4
0.2
0
FRA−VEY−10
FRA−VEY−3
FRA−VEY−8
FRA−VEY−9
FRA−VEY−1
FRA−VEY−14
FRA−VEY−16
FRA−VEY−5
FRA−VEY−11
FRA−VEY−12
FRA−VEY−17
FRA−VEY−6
FRA−VEY−15
FRA−VEY−7
FRA−VEY−13
FRA−VEY−2

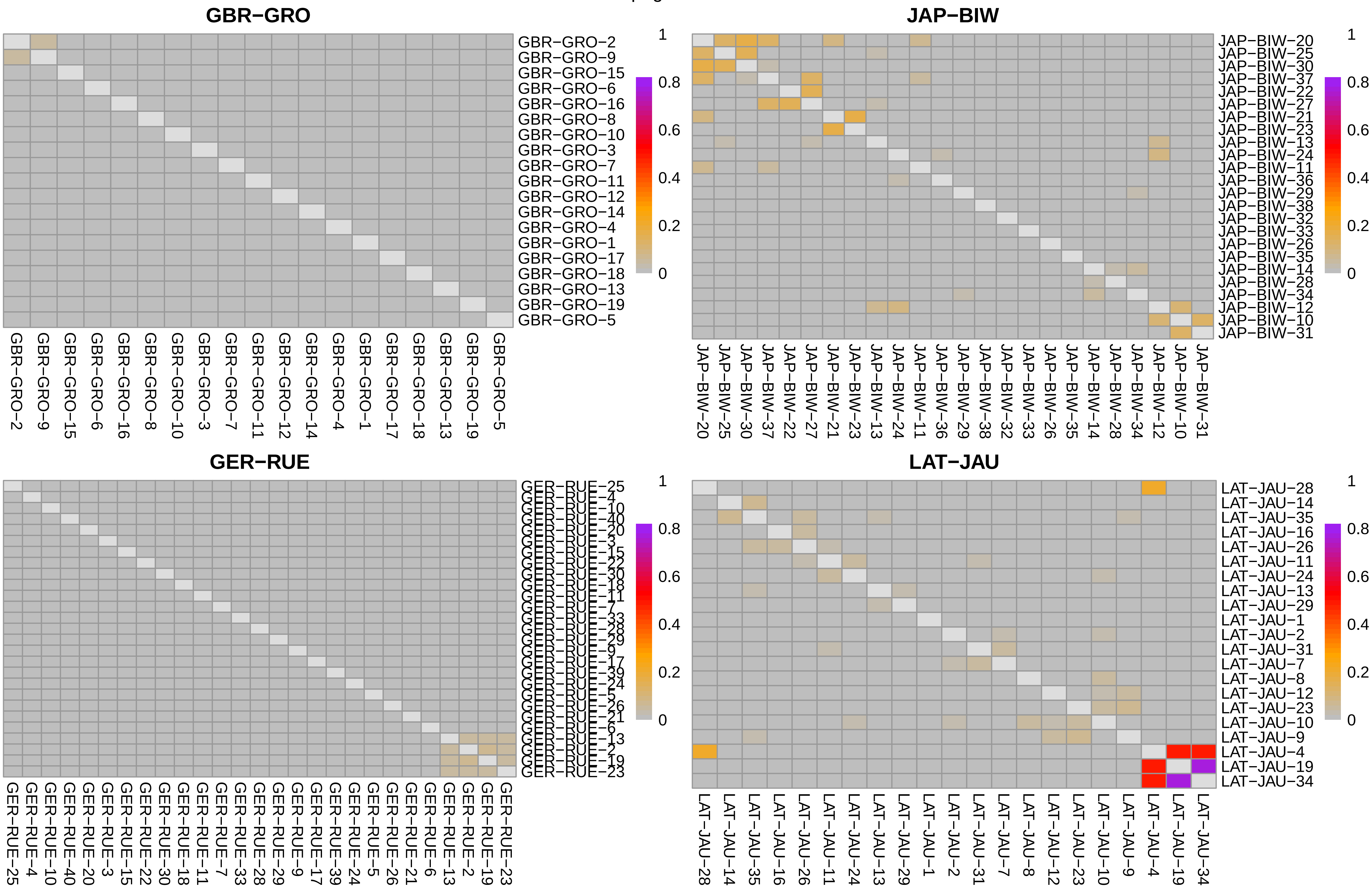
GBR−GRO
GBR−GRO−2 GBR−GRO−9 GBR−GRO−15 GBR−GRO−6 GBR−GRO−16 GBR−GRO−8 GBR−GRO−10 GBR−GRO−3 GBR−GRO−7 GBR−GRO−11 GBR−GRO−12 GBR−GRO−14 GBR−GRO−4 GBR−GRO−1 GBR−GRO−17 GBR−GRO−18 GBR−GRO−13 GBR−GRO−19 GBR−GRO−5
1 0.8 0.6 0.4 0.2 0
JAP−BIW
JAP−BIW−20 JAP−BIW−25 JAP−BIW−30 JAP−BIW−37 JAP−BIW−22 JAP−BIW−27 JAP−BIW−21 JAP−BIW−23 JAP−BIW−13 JAP−BIW−24 JAP−BIW−11 JAP−BIW−36 JAP−BIW−29 JAP−BIW−38 JAP−BIW−32 JAP−BIW−33 JAP−BIW−26 JAP−BIW−35 JAP−BIW−14 JAP−BIW−28 JAP−BIW−34 JAP−BIW−12 JAP−BIW−10 JAP−BIW−31
1 0.8 0.6 0.4 0.2 0
GER−RUE
GER−RUE−25 GER−RUE−4 GER−RUE−10 GER−RUE−40 GER−RUE−20 GER−RUE−3 GER−RUE−15 GER−RUE−22 GER−RUE−30 GER−RUE−18 GER−RUE−11 GER−RUE−7 GER−RUE−33 GER−RUE−28 GER−RUE−29 GER−RUE−9 GER−RUE−17 GER−RUE−39 GER−RUE−24 GER−RUE−5 GER−RUE−26 GER−RUE−21 GER−RUE−6 GER−RUE−13 GER−RUE−2 GER−RUE−19 GER−RUE−23
1 0.8 0.6 0.4 0.2 0
LAT−JAU
LAT−JAU−28 LAT−JAU−14 LAT−JAU−35 LAT−JAU−16 LAT−JAU−26 LAT−JAU−11 LAT−JAU−24 LAT−JAU−13 LAT−JAU−29 LAT−JAU−1 LAT−JAU−2 LAT−JAU−31 LAT−JAU−7 LAT−JAU−8 LAT−JAU−12 LAT−JAU−23 LAT−JAU−10 LAT−JAU−9 LAT−JAU−4 LAT−JAU−19 LAT−JAU−34
1 0.8 0.6 0.4 0.2 0

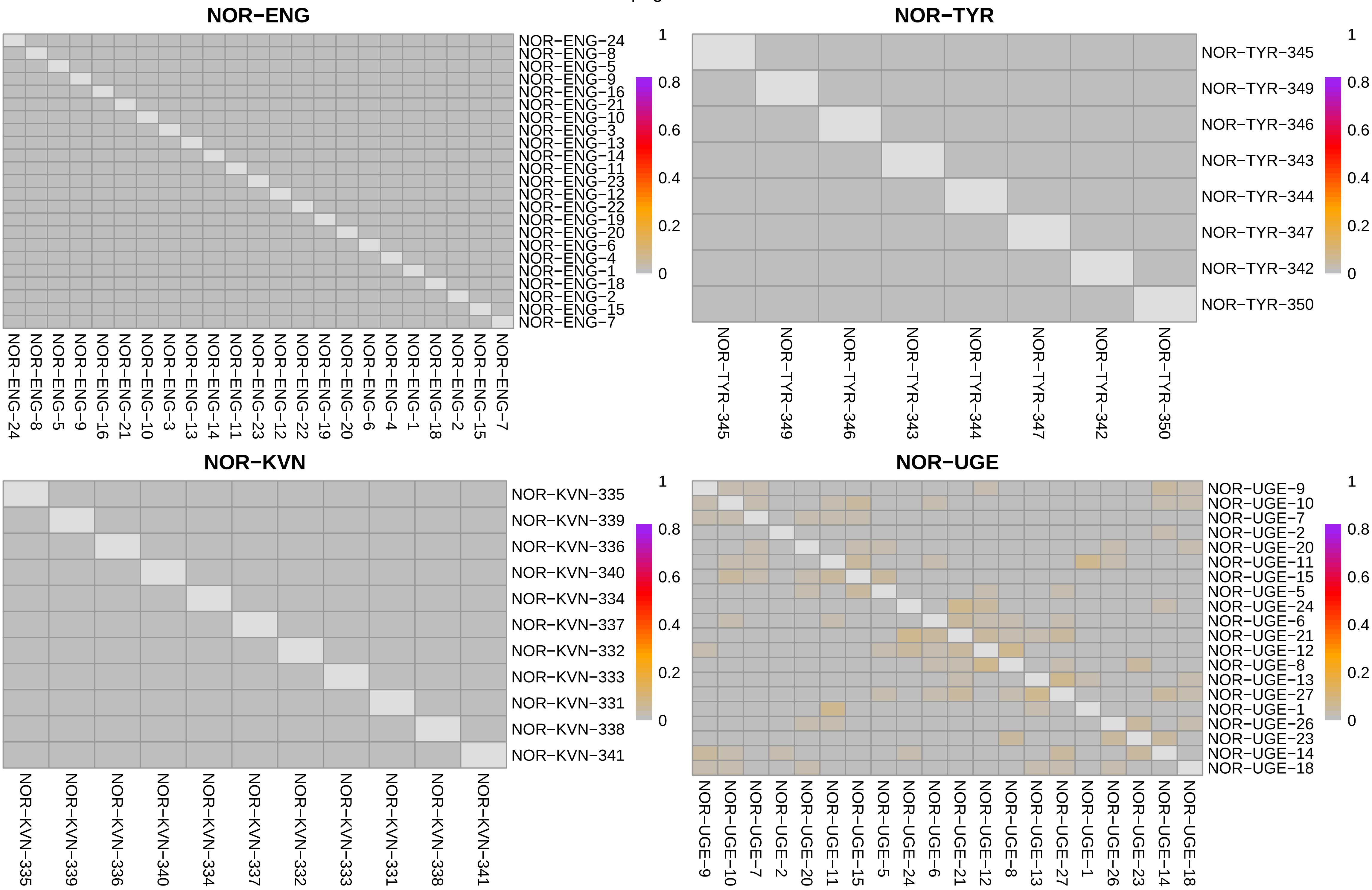
NOR−ENG
NOR−ENG−24
NOR−ENG−8
NOR−ENG−5
NOR−ENG−9
NOR−ENG−16
NOR−ENG−21
NOR−ENG−10
NOR−ENG−3
NOR−ENG−13
NOR−ENG−14
NOR−ENG−11
NOR−ENG−23
NOR−ENG−12
NOR−ENG−22
NOR−ENG−19
NOR−ENG−20
NOR−ENG−6
NOR−ENG−4
NOR−ENG−1
NOR−ENG−18
NOR−ENG−2
NOR−ENG−15
NOR−ENG−7
1
0.8
0.6
0.4
0.2
0
NOR−TYR
NOR−TYR−345
NOR−TYR−349
NOR−TYR−346
NOR−TYR−343
NOR−TYR−344
NOR−TYR−347
NOR−TYR−342
NOR−TYR−350
NOR−KVN
NOR−KVN−335
NOR−KVN−339
NOR−KVN−336
NOR−KVN−340
NOR−KVN−334
NOR−KVN−337
NOR−KVN−332
NOR−KVN−333
NOR−KVN−331
NOR−KVN−338
NOR−KVN−341
NOR−UGE
NOR−UGE−9
NOR−UGE−10
NOR−UGE−7
NOR−UGE−2
NOR−UGE−20
NOR−UGE−11
NOR−UGE−15
NOR−UGE−5
NOR−UGE−24
NOR−UGE−6
NOR−UGE−21
NOR−UGE−12
NOR−UGE−8
NOR−UGE−13
NOR−UGE−27
NOR−UGE−1
NOR−UGE−26
NOR−UGE−23
NOR−UGE−14
NOR−UGE−18

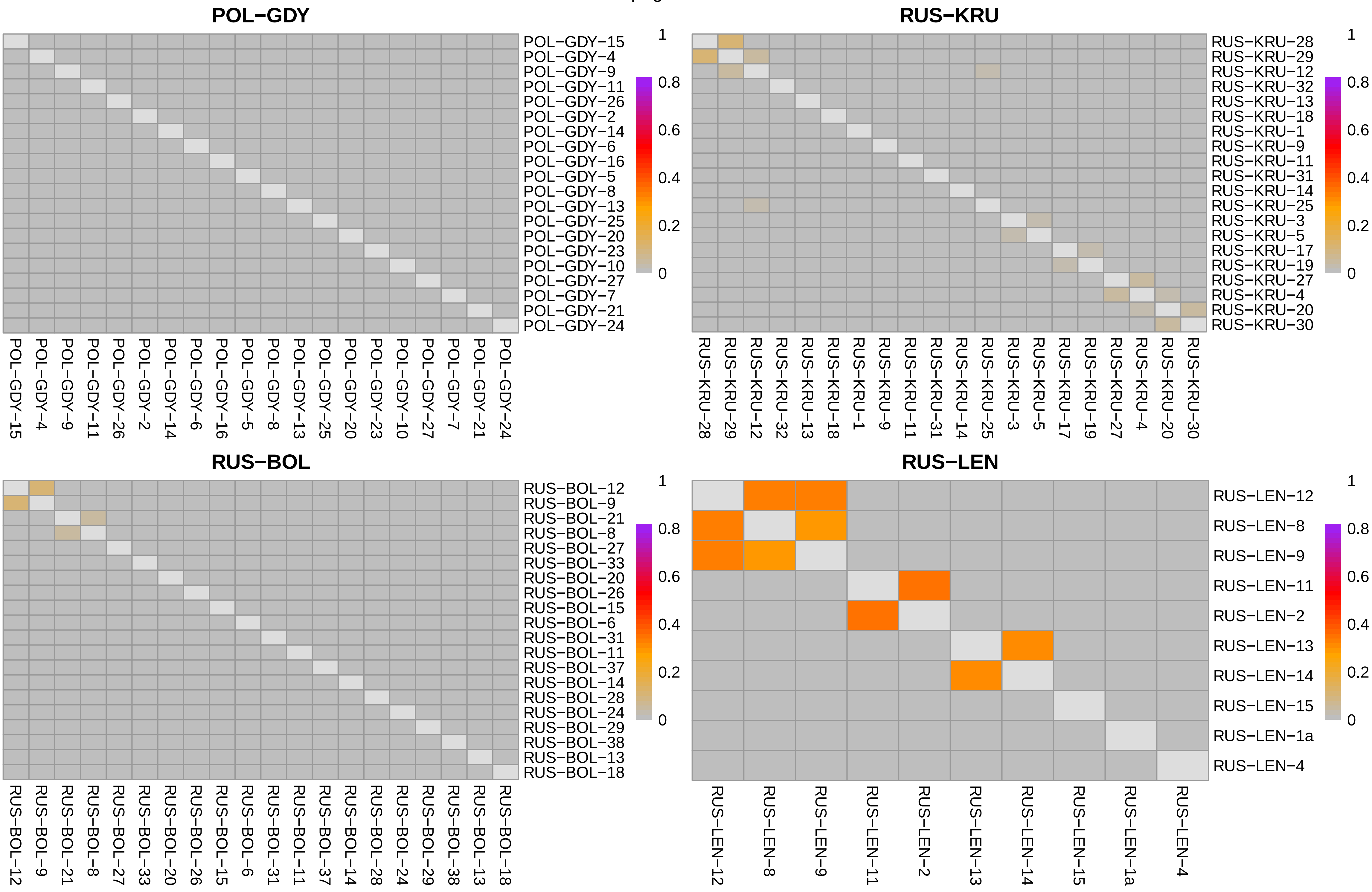
POL−GDY
POL−GDY−15
POL−GDY−4
POL−GDY−9
POL−GDY−11
POL−GDY−26
POL−GDY−2
POL−GDY−14
POL−GDY−6
POL−GDY−16
POL−GDY−5
POL−GDY−8
POL−GDY−13
POL−GDY−25
POL−GDY−20
POL−GDY−23
POL−GDY−10
POL−GDY−27
POL−GDY−7
POL−GDY−21
POL−GDY−24
1
0.8
0.6
0.4
0.2
0
RUS−KRU
RUS−KRU−28
RUS−KRU−29
RUS−KRU−12
RUS−KRU−32
RUS−KRU−13
RUS−KRU−18
RUS−KRU−1
RUS−KRU−9
RUS−KRU−11
RUS−KRU−31
RUS−KRU−14
RUS−KRU−25
RUS−KRU−3
RUS−KRU−5
RUS−KRU−17
RUS−KRU−19
RUS−KRU−27
RUS−KRU−4
RUS−KRU−20
RUS−KRU−30
RUS−BOL
RUS−BOL−12
RUS−BOL−9
RUS−BOL−21
RUS−BOL−8
RUS−BOL−27
RUS−BOL−33
RUS−BOL−20
RUS−BOL−26
RUS−BOL−15
RUS−BOL−6
RUS−BOL−31
RUS−BOL−11
RUS−BOL−37
RUS−BOL−14
RUS−BOL−28
RUS−BOL−24
RUS−BOL−29
RUS−BOL−38
RUS−BOL−13
RUS−BOL−18
RUS−LEN
RUS−LEN−12
RUS−LEN−8
RUS−LEN−9
RUS−LEN−11
RUS−LEN−2
RUS−LEN−13
RUS−LEN−14
RUS−LEN−15
RUS−LEN−1a
RUS−LEN−4

### RUS−LEV

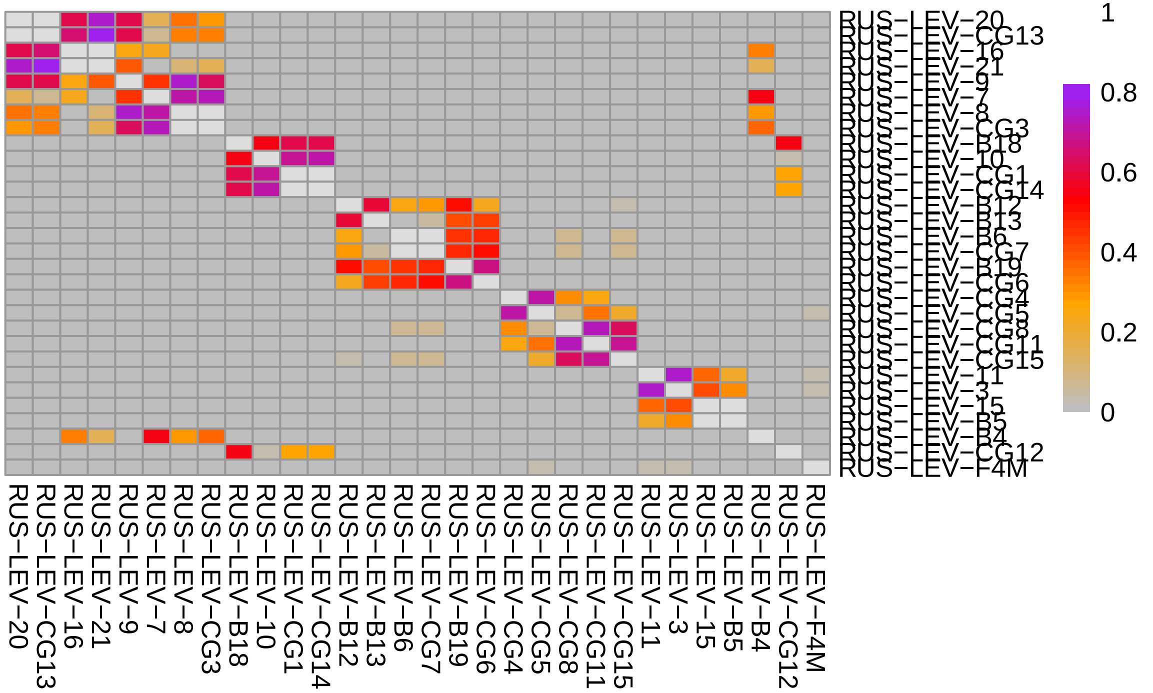


### SCO−HAR

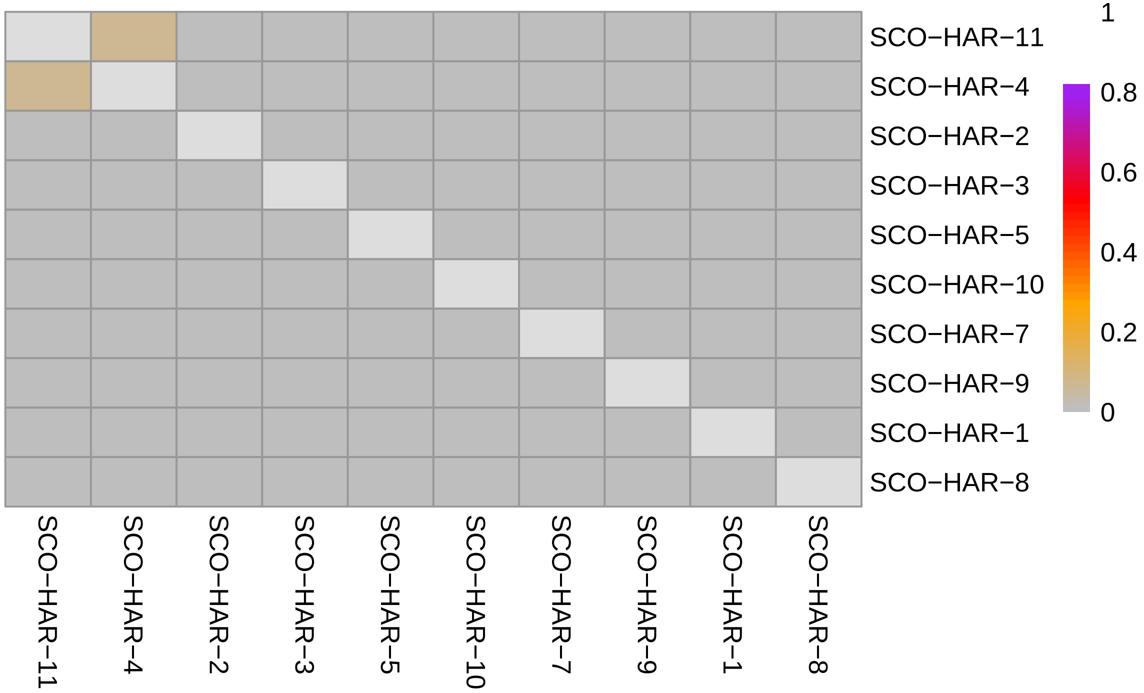


### RUS−MAS

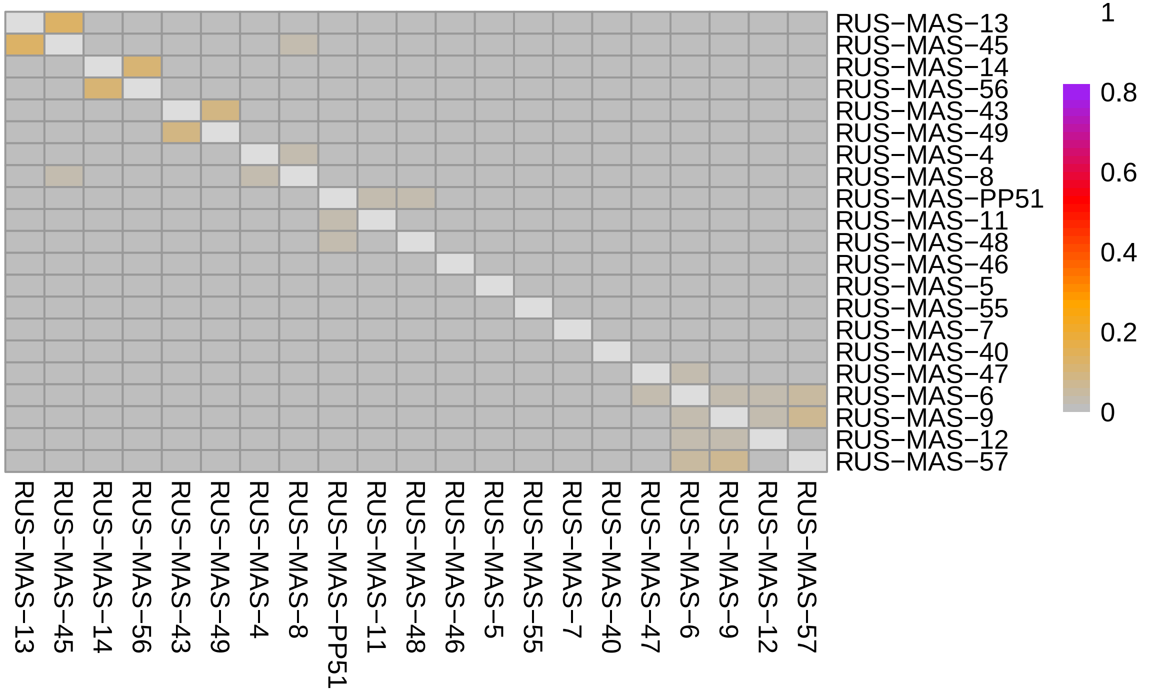


### SWE−ABB

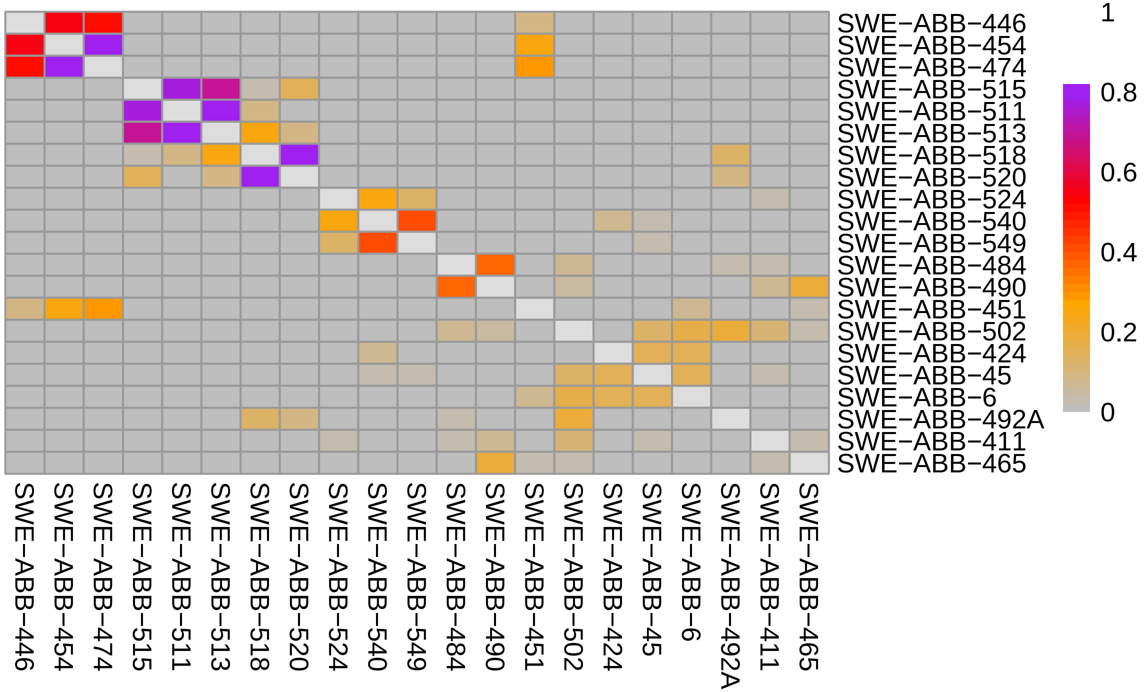

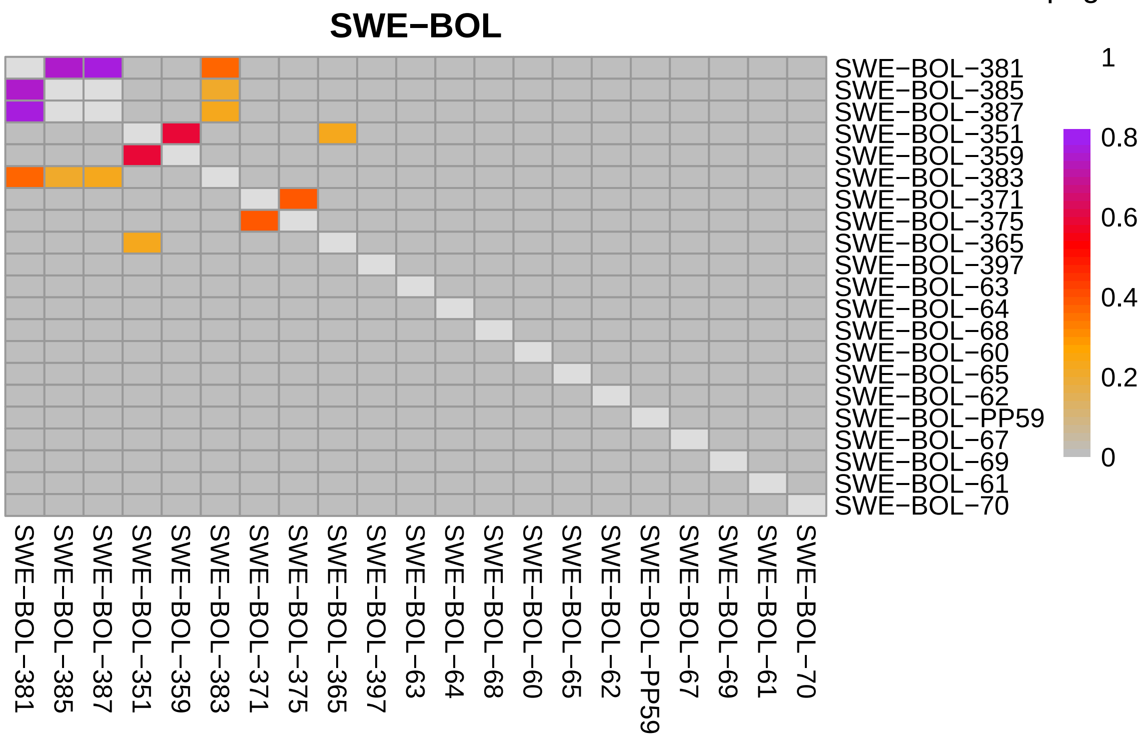
SWE−BOL
1
0.8
0.6
0.4
0.2
0
SWE−BOL−381
SWE−BOL−385
SWE−BOL−387
SWE−BOL−351
SWE−BOL−359
SWE−BOL−383
SWE−BOL−371
SWE−BOL−375
SWE−BOL−365
SWE−BOL−397
SWE−BOL−63
SWE−BOL−64
SWE−BOL−68
SWE−BOL−60
SWE−BOL−65
SWE−BOL−62
SWE−BOL−PP59
SWE−BOL−67
SWE−BOL−69
SWE−BOL−61
SWE−BOL−70

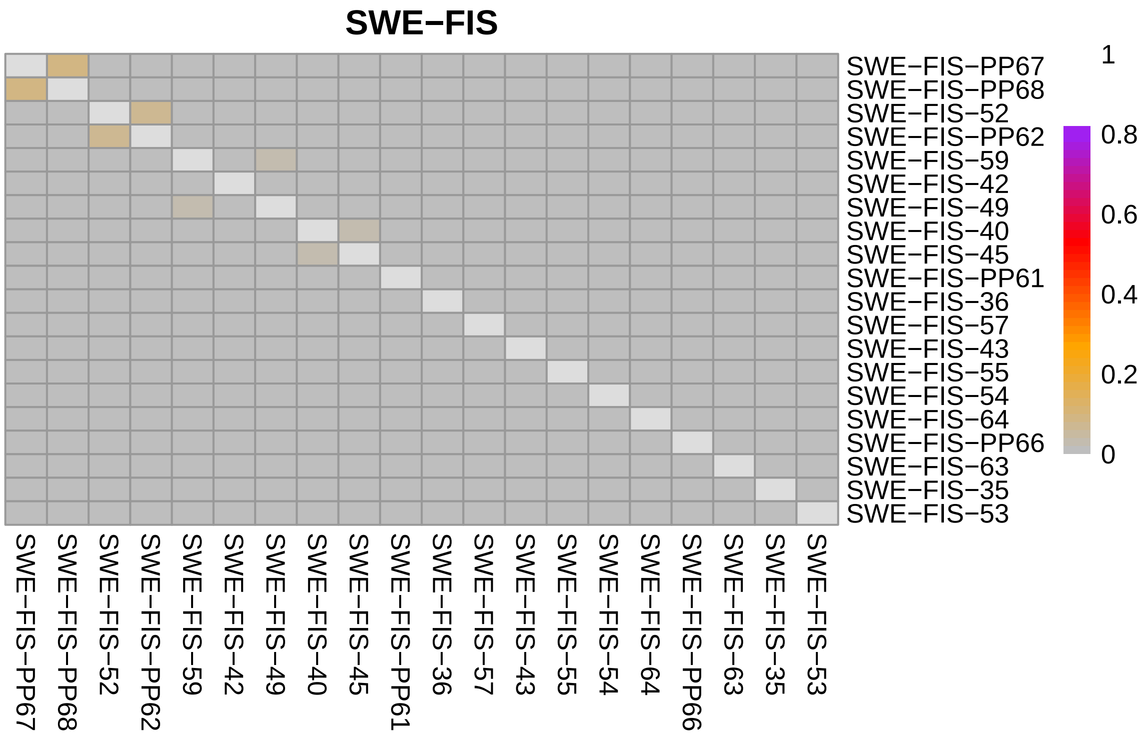
SWE−FIS
1
0.8
0.6
0.4
0.2
0
SWE−FIS−PP67
SWE−FIS−PP68
SWE−FIS−52
SWE−FIS−PP62
SWE−FIS−59
SWE−FIS−42
SWE−FIS−49
SWE−FIS−40
SWE−FIS−45
SWE−FIS−PP61
SWE−FIS−36
SWE−FIS−57
SWE−FIS−43
SWE−FIS−55
SWE−FIS−54
SWE−FIS−64
SWE−FIS−PP66
SWE−FIS−63
SWE−FIS−35
SWE−FIS−53

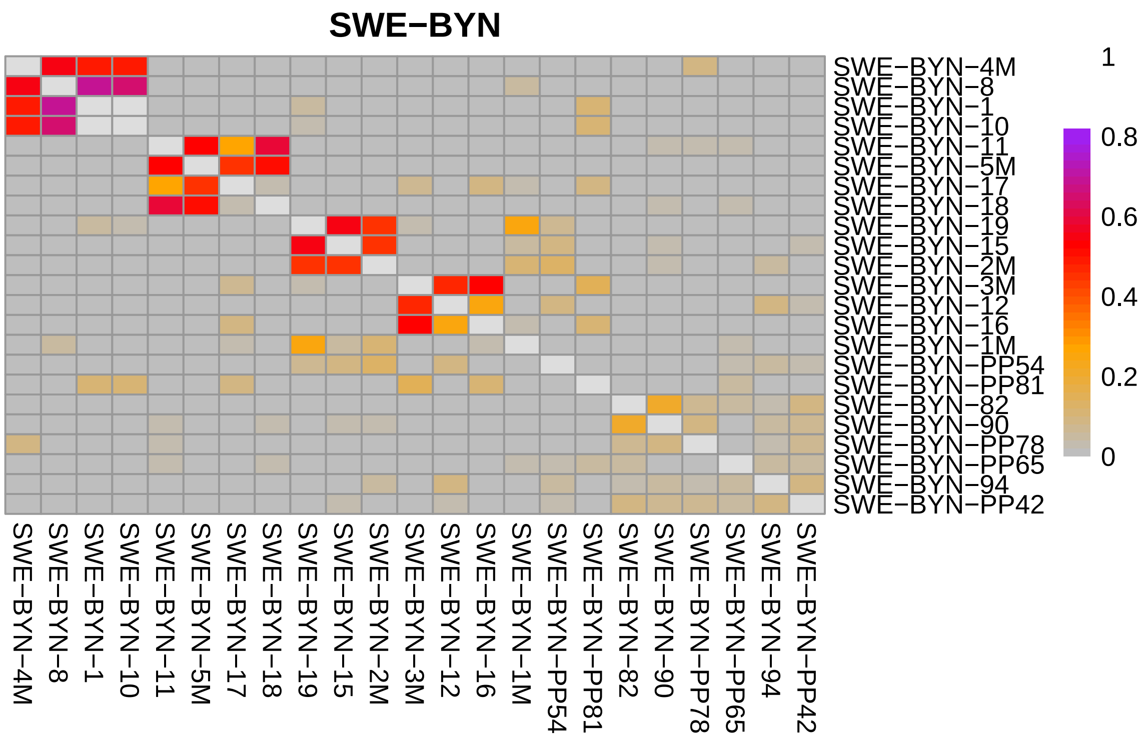
SWE−BYN
1
0.8
0.6
0.4
0.2
0
SWE−BYN−4M
SWE−BYN−8
SWE−BYN−1
SWE−BYN−10
SWE−BYN−11
SWE−BYN−5M
SWE−BYN−17
SWE−BYN−18
SWE−BYN−19
SWE−BYN−15
SWE−BYN−2M
SWE−BYN−3M
SWE−BYN−12
SWE−BYN−16
SWE−BYN−1M
SWE−BYN−PP54
SWE−BYN−PP81
SWE−BYN−82
SWE−BYN−90
SWE−BYN−PP78
SWE−BYN−PP65
SWE−BYN−94
SWE−BYN−PP42

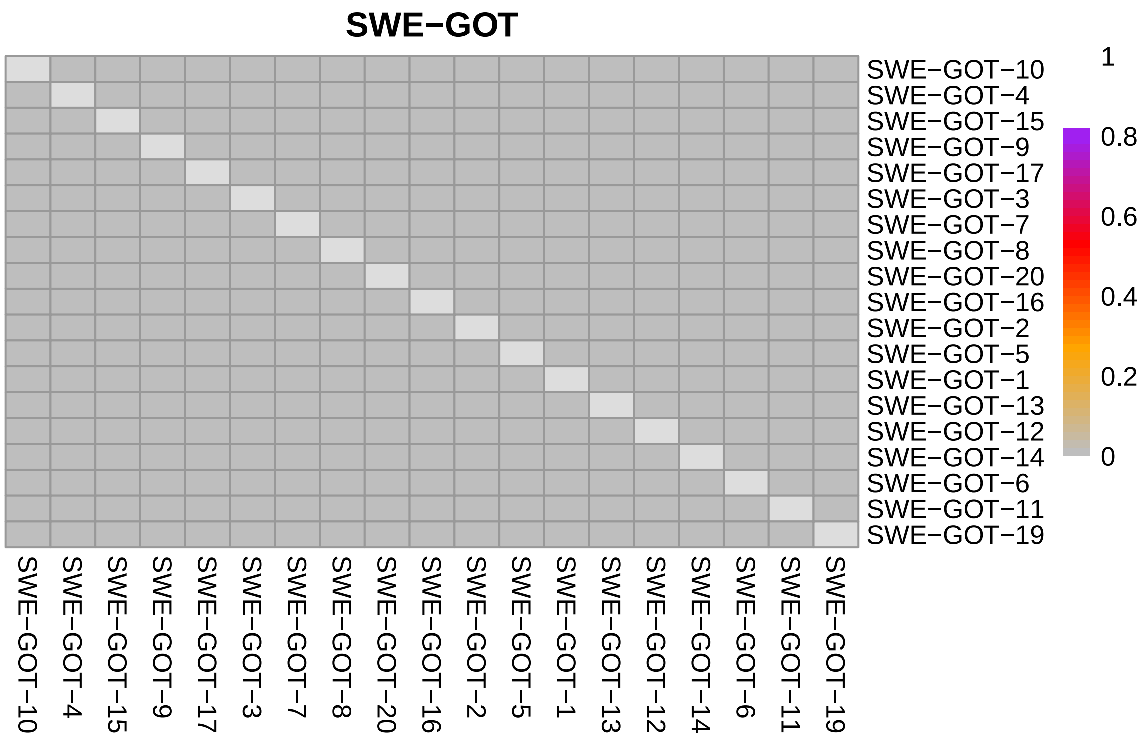
SWE−GOT
1
0.8
0.6
0.4
0.2
0
SWE−GOT−10
SWE−GOT−4
SWE−GOT−15
SWE−GOT−9
SWE−GOT−17
SWE−GOT−3
SWE−GOT−7
SWE−GOT−8
SWE−GOT−20
SWE−GOT−16
SWE−GOT−2
SWE−GOT−5
SWE−GOT−1
SWE−GOT−13
SWE−GOT−12
SWE−GOT−14
SWE−GOT−6
SWE−GOT−11
SWE−GOT−19

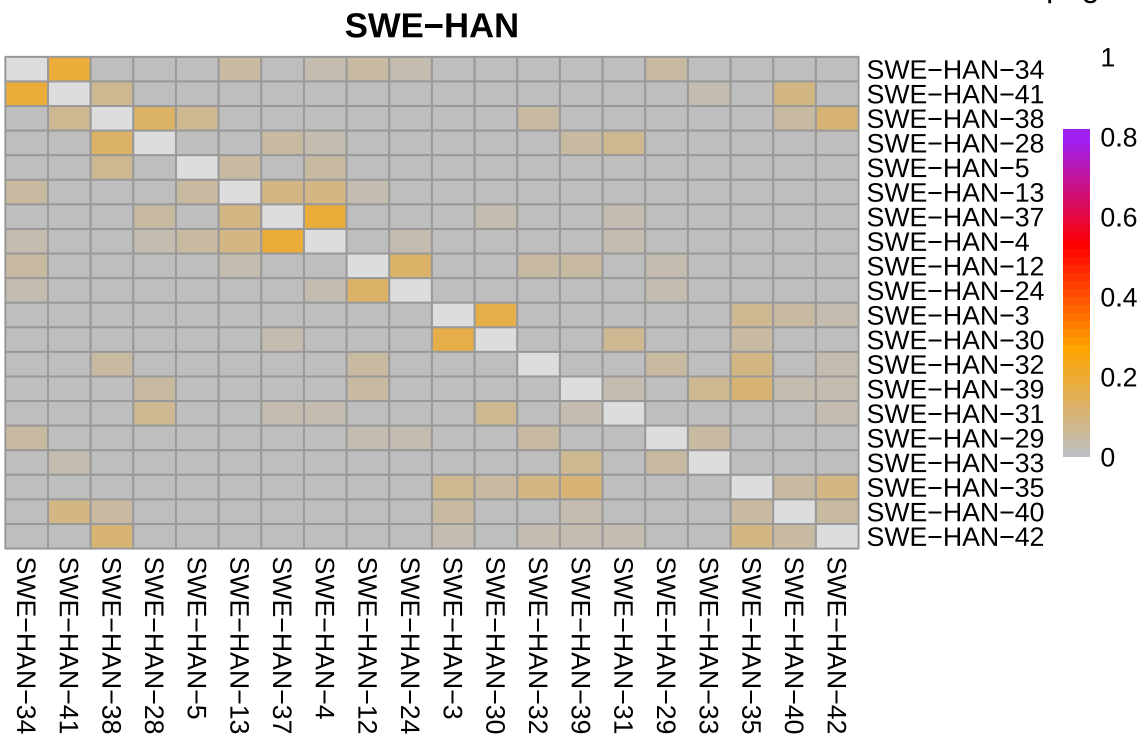
SWE−HAN
SWE−HAN−34
SWE−HAN−41
SWE−HAN−38
SWE−HAN−28
SWE−HAN−5
SWE−HAN−13
SWE−HAN−37
SWE−HAN−4
SWE−HAN−12
SWE−HAN−24
SWE−HAN−3
SWE−HAN−30
SWE−HAN−32
SWE−HAN−39
SWE−HAN−31
SWE−HAN−29
SWE−HAN−33
SWE−HAN−35
SWE−HAN−40
SWE−HAN−42
1
0.8
0.6
0.4
0.2
0

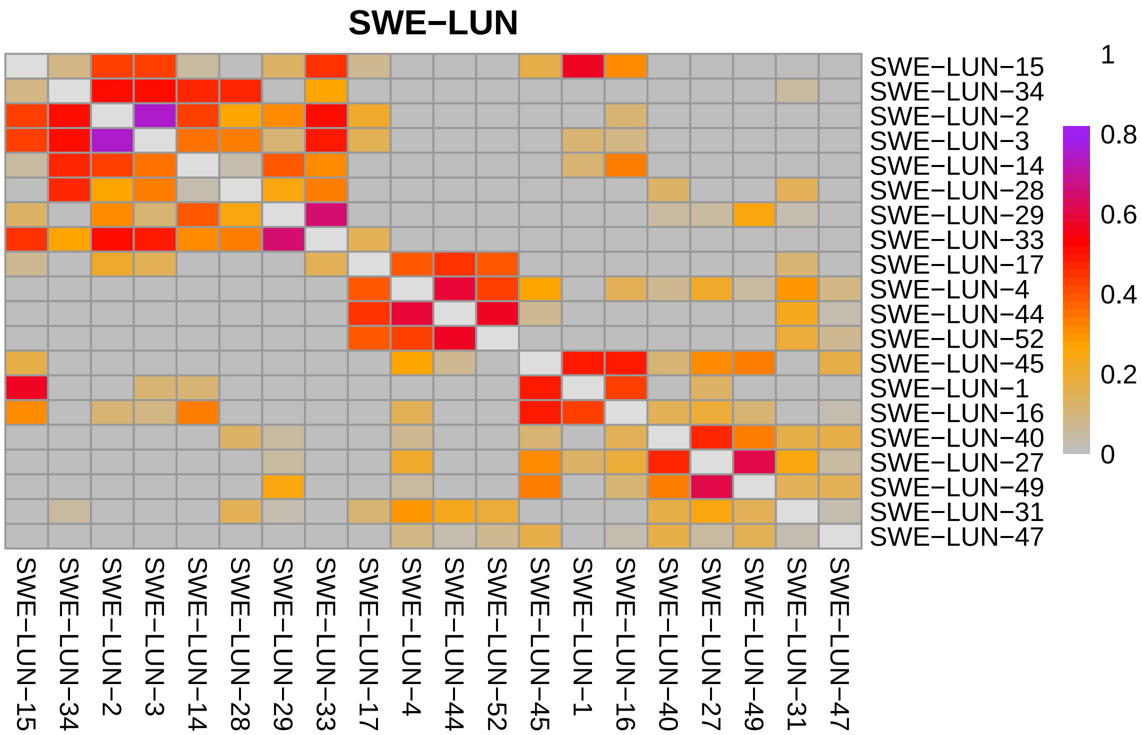
SWE−LUN
SWE−LUN−15
SWE−LUN−34
SWE−LUN−2
SWE−LUN−3
SWE−LUN−14
SWE−LUN−28
SWE−LUN−29
SWE−LUN−33
SWE−LUN−17
SWE−LUN−4
SWE−LUN−44
SWE−LUN−52
SWE−LUN−45
SWE−LUN−1
SWE−LUN−16
SWE−LUN−40
SWE−LUN−27
SWE−LUN−49
SWE−LUN−31
SWE−LUN−47
1
0.8
0.6
0.4
0.2
0

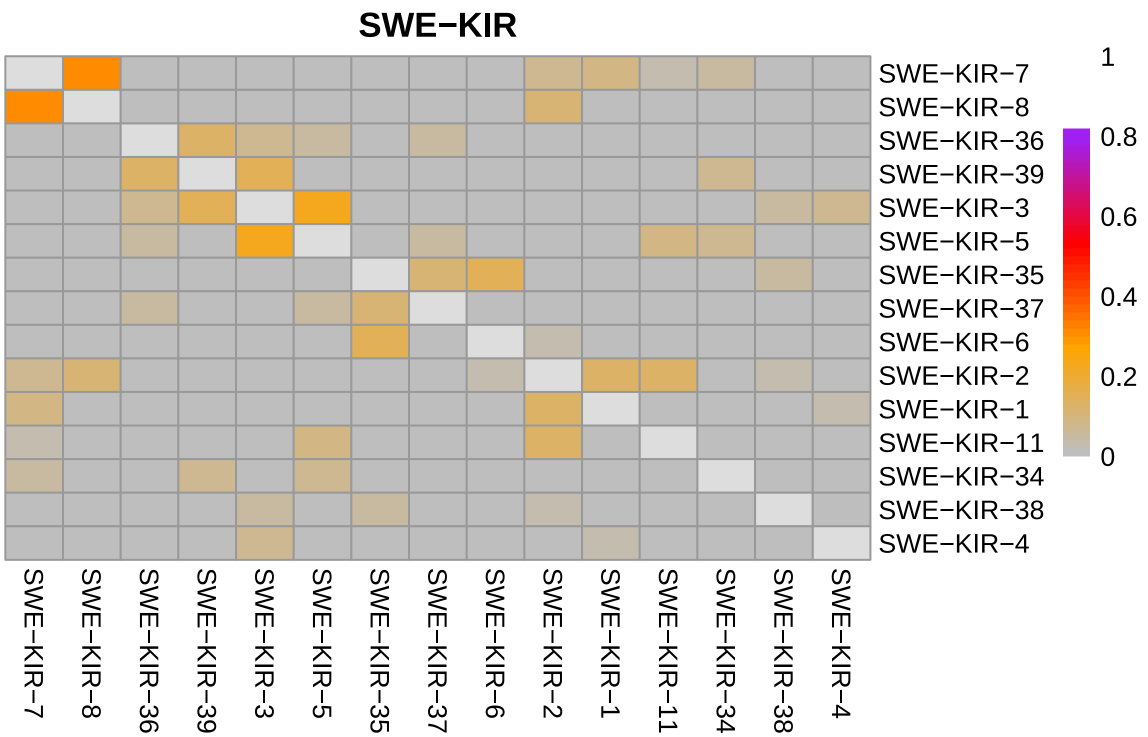
SWE−KIR
SWE−KIR−7
SWE−KIR−8
SWE−KIR−36
SWE−KIR−39
SWE−KIR−3
SWE−KIR−5
SWE−KIR−35
SWE−KIR−37
SWE−KIR−6
SWE−KIR−2
SWE−KIR−1
SWE−KIR−11
SWE−KIR−34
SWE−KIR−38
SWE−KIR−4
1
0.8
0.6
0.4
0.2
0

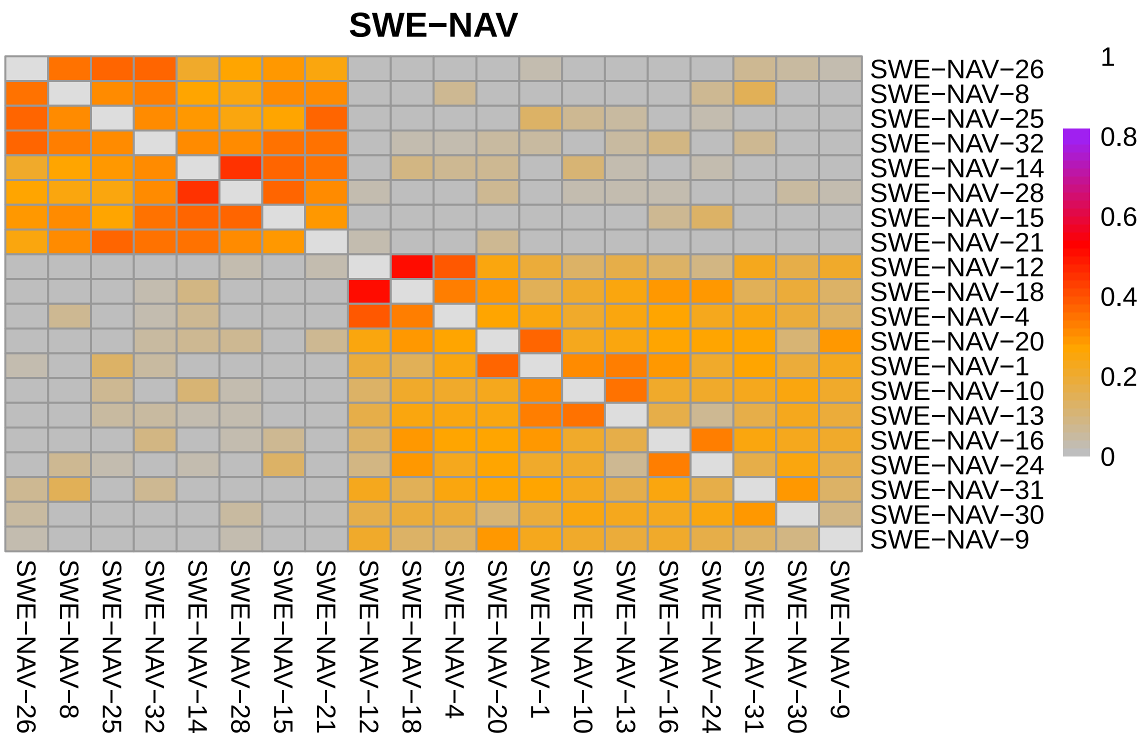
SWE−NAV
SWE−NAV−26
SWE−NAV−8
SWE−NAV−25
SWE−NAV−32
SWE−NAV−14
SWE−NAV−28
SWE−NAV−15
SWE−NAV−21
SWE−NAV−12
SWE−NAV−18
SWE−NAV−4
SWE−NAV−20
SWE−NAV−1
SWE−NAV−10
SWE−NAV−13
SWE−NAV−16
SWE−NAV−24
SWE−NAV−31
SWE−NAV−30
SWE−NAV−9
1
0.8
0.6
0.4
0.2
0

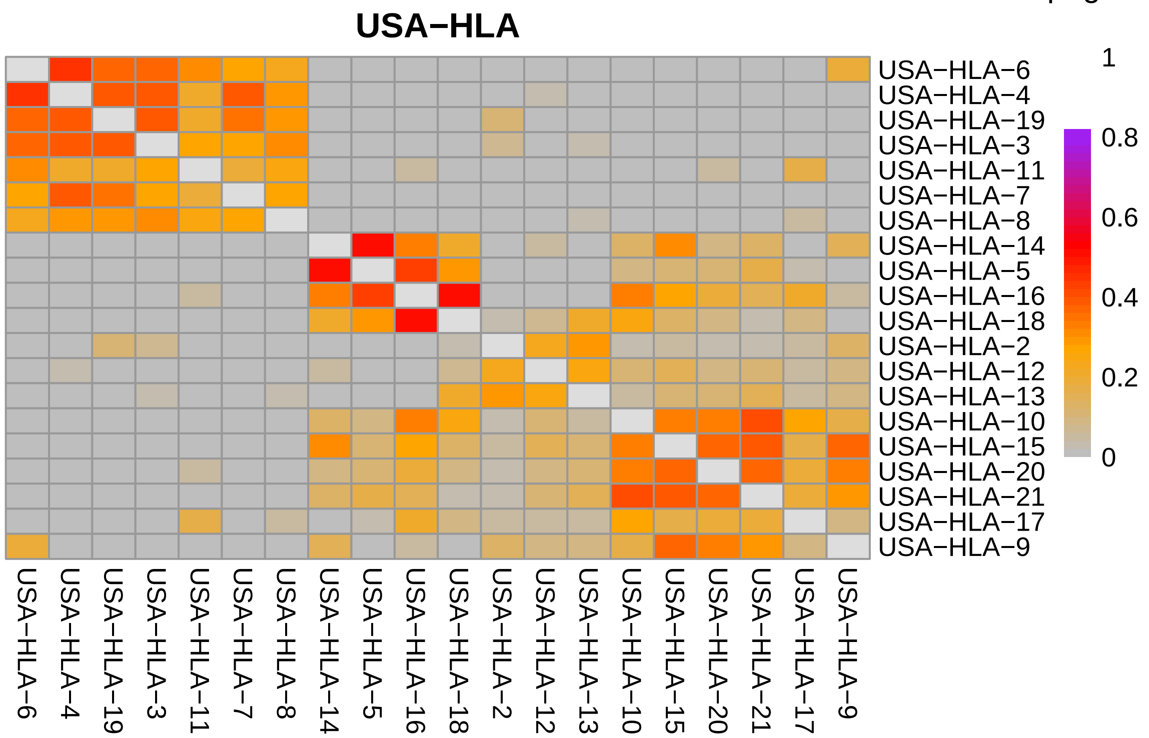
USA−HLA
USA−HLA−6
USA−HLA−4
USA−HLA−19
USA−HLA−3
USA−HLA−11
USA−HLA−7
USA−HLA−8
USA−HLA−14
USA−HLA−5
USA−HLA−16
USA−HLA−18
USA−HLA−2
USA−HLA−12
USA−HLA−13
USA−HLA−10
USA−HLA−15
USA−HLA−20
USA−HLA−21
USA−HLA−17
USA−HLA−9
1
0.8
0.6
0.4
0.2
0
