## Supplemental figs and tables for "Estimating recent and historical effective population size of marine and freshwater sticklebacks": Supplemental figure legends.pdf

### Supplementary Figure Legend

Figure S1. Map illustrating sampling locations of the studied populations, derived from Feng et al., (2022). The ecotypes are indicated by the colors on top.

Figure S2. Linkage disequilibrium (LD) decay curve in each population.

Figure S3. Relatedness ($r_{xy}$) for pairs of individuals within each population. Colors representing specific relatedness classes.

Figure S4. Effective population size ($N_e$) of each population estimated with GONE. The x-axis shows the time in generations before the sampling. Within each panel, the solid and dashed line correspond to the result of using a recombination fraction of 0.01 and 0.05, respectively.

Figure S5. Estimates of contemporary effective population size ($N_e^C$) for 45 nine-spined stickleback populations. a). $N_e^C$ and its 90% confidence intervals estimated using CurrentNe2 and GONE (with *hc* = 0.01 and 0.05). Populations are arranged in the same order as in Fig. 1 and the background color indicates the ecotype. b). Populations inferred to have experienced a drastic decline in $N_e$ in GONE analysis. For exact $N_e^C$ values, see Table S5.

Figure S6. Historical effective population sizes ($N_e$) for each population inferred with MSMC2. The x-axis shows the time in years before present based on generation time of two years and mutation rate of $4.37\times10^{-9}$ per base pair per generation. The grey, blue and orange bars at the bottom indicate the times for the formation of the Baltic Sea, the Last Glacial Period and the Last Glacial Maximum, respectively. Within each panel, the thin lines correspond to the original inferences and 20 rounds of bootstrap replicates, and the bold lines show the original inferences of each population.

Figure S7. Historical effective population sizes ($N_e$) for populations within different ecotypes inferred with MSMC2. a) Marine populations. b) Coastal and freshwater populations. c) Lake and stream populations. d) Pond populations. The x-axis shows the time in years before present based on generation time of two years and mutation rate of 4.37×10-9 per base pair per generation. The grey, blue and orange bars at the bottom indicate the times for the formation of the Baltic Sea, the Last Glacial Period and the Last Glacial Maximum, respectively. Within each panel, the thin lines correspond to the original inferences and 20 rounds of bootstrap replicates, and the bold lines show representative populations from each ecotype.

Figure S8. Temporal reconstruction of effective population size ($N_e$) for 45 nine-spined stickleback populations over the past 30,000 years. a) The colors represent $N_e$ estimates at key time points with the scale given at the top. The estimates for the last 200 years were derived with GONE (*hc* = 0.05), while those for 4,000 to 30,000 years ago were obtained with MSMC2. Asterisks indicate estimates influenced by population structure as inferred by manual inspection following the patterns in Fig. 2F of Santiago et al. (2020). Long-term $N_e$ values were calculated from genetic diversity ($\pi$). Key time points were chosen to reflect significant geographic events: 4,000 years ago marks the onset of the current Baltic Sea stage, 12,600 years ago the start of the Baltic Ice Lake stage, and 21,000-27,000 years ago corresponds to the Last Glacial Maximum. The color next to population labels represents the ecotypes. b) The demographic history of a representative population, FIN-PYO, over the past 80,000 years. The dashed lines connect the time points in panels a) and b).
