## Supplementary figures and images for "Estimating recent and historical effective population size of marine and freshwater sticklebacks"

### FigS1.pdf

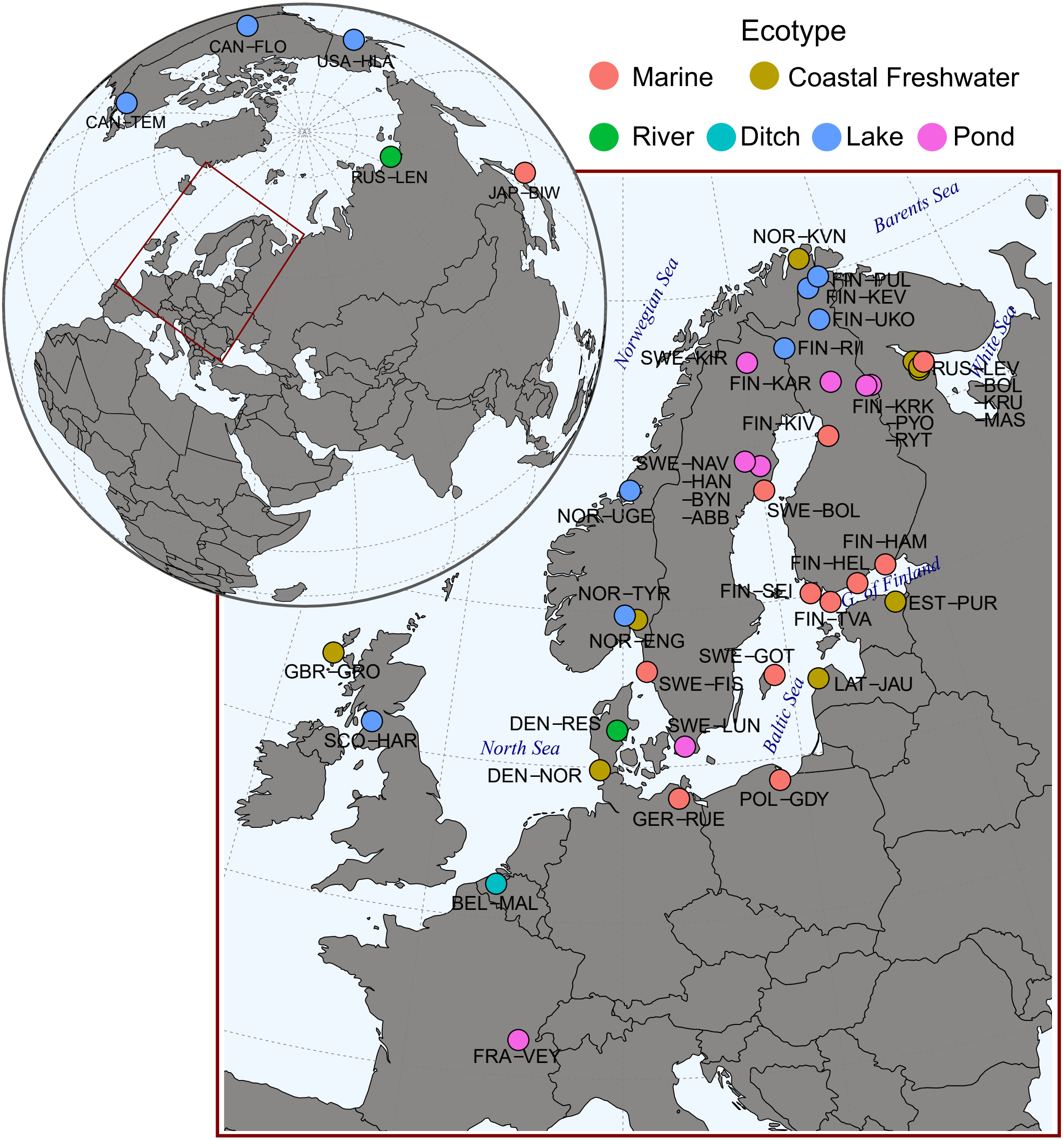

### FigS2.pdf

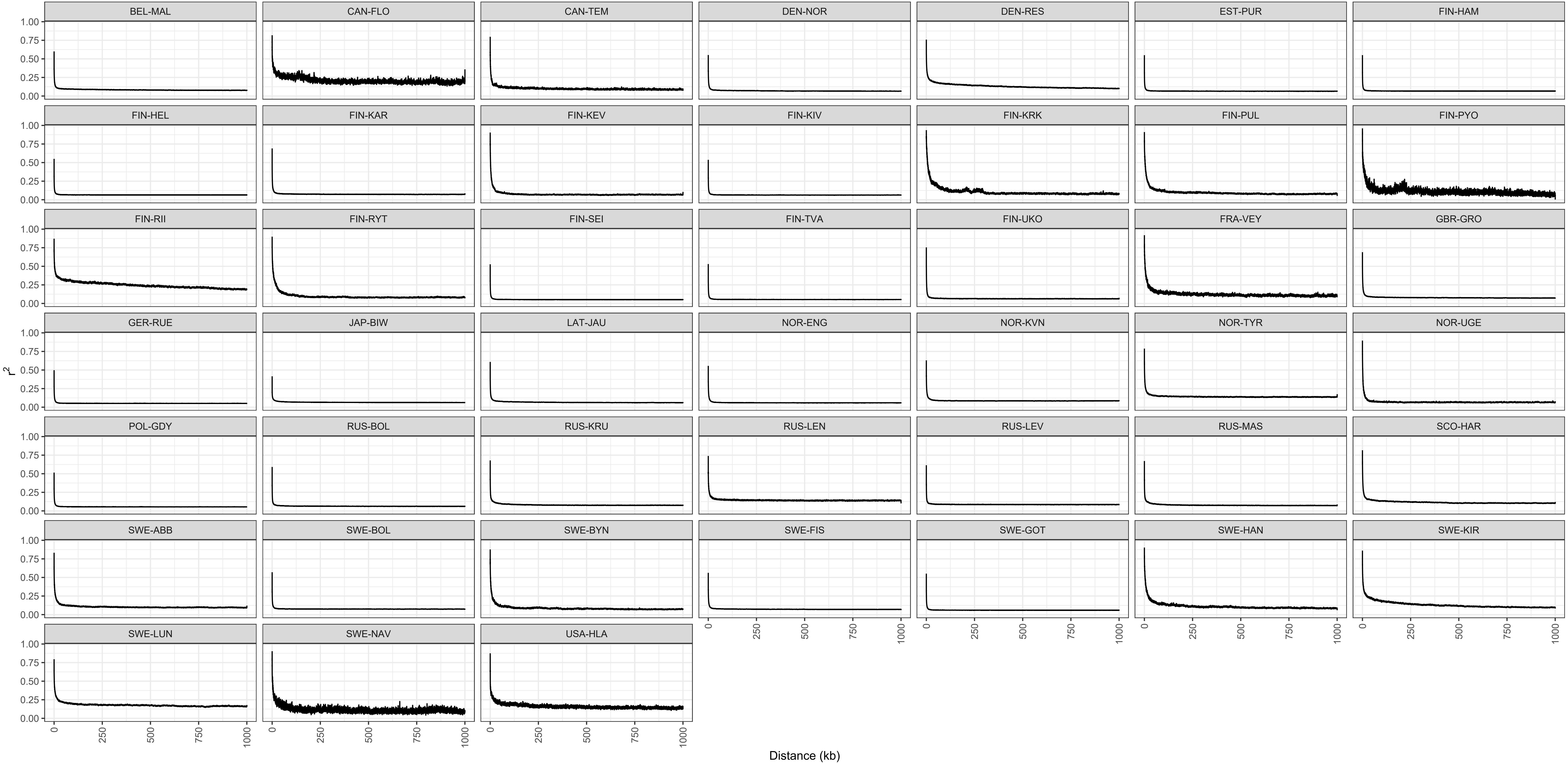

### FigS4.pdf

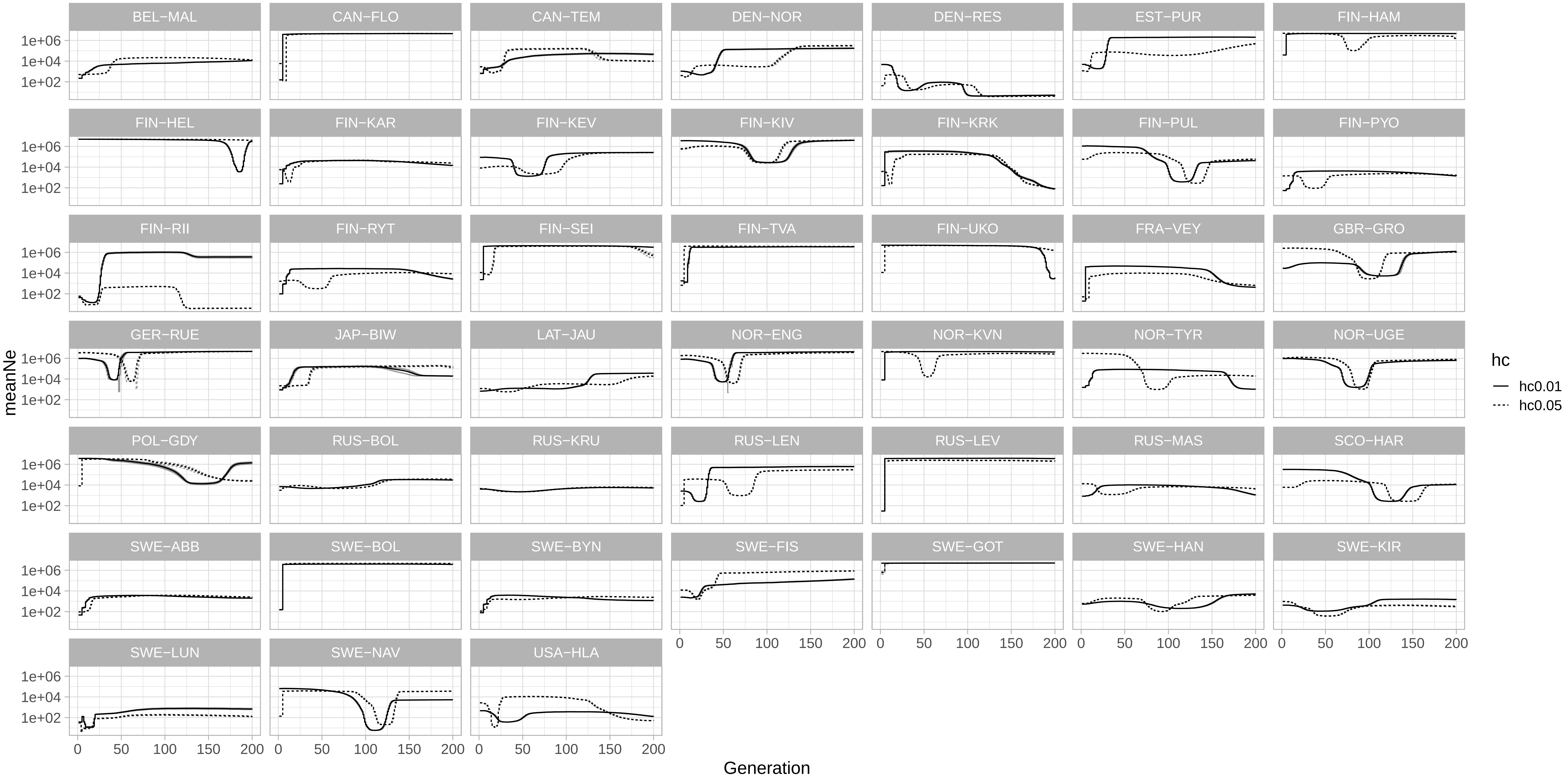

### FigS5.pdf

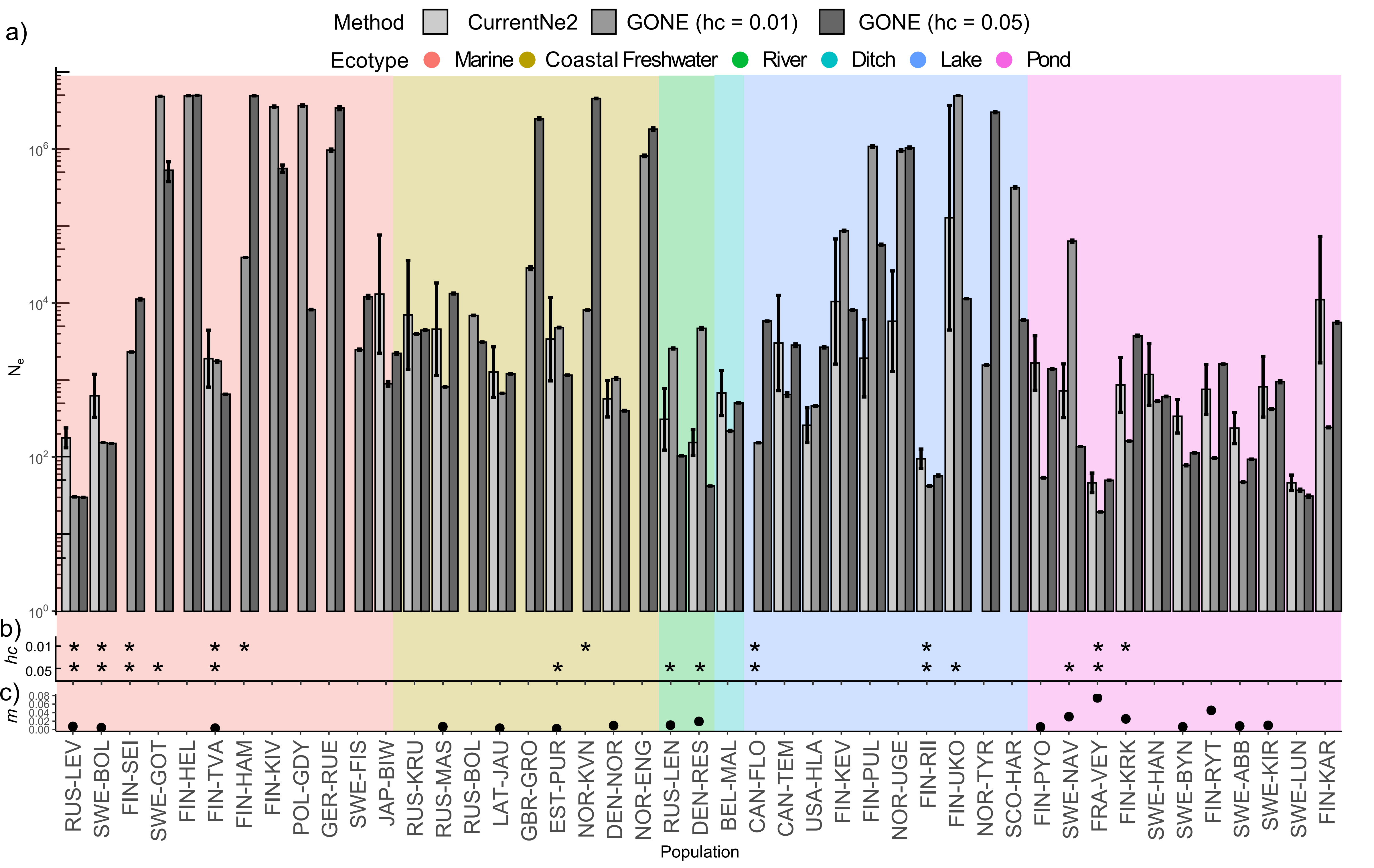

### FigS6.pdf

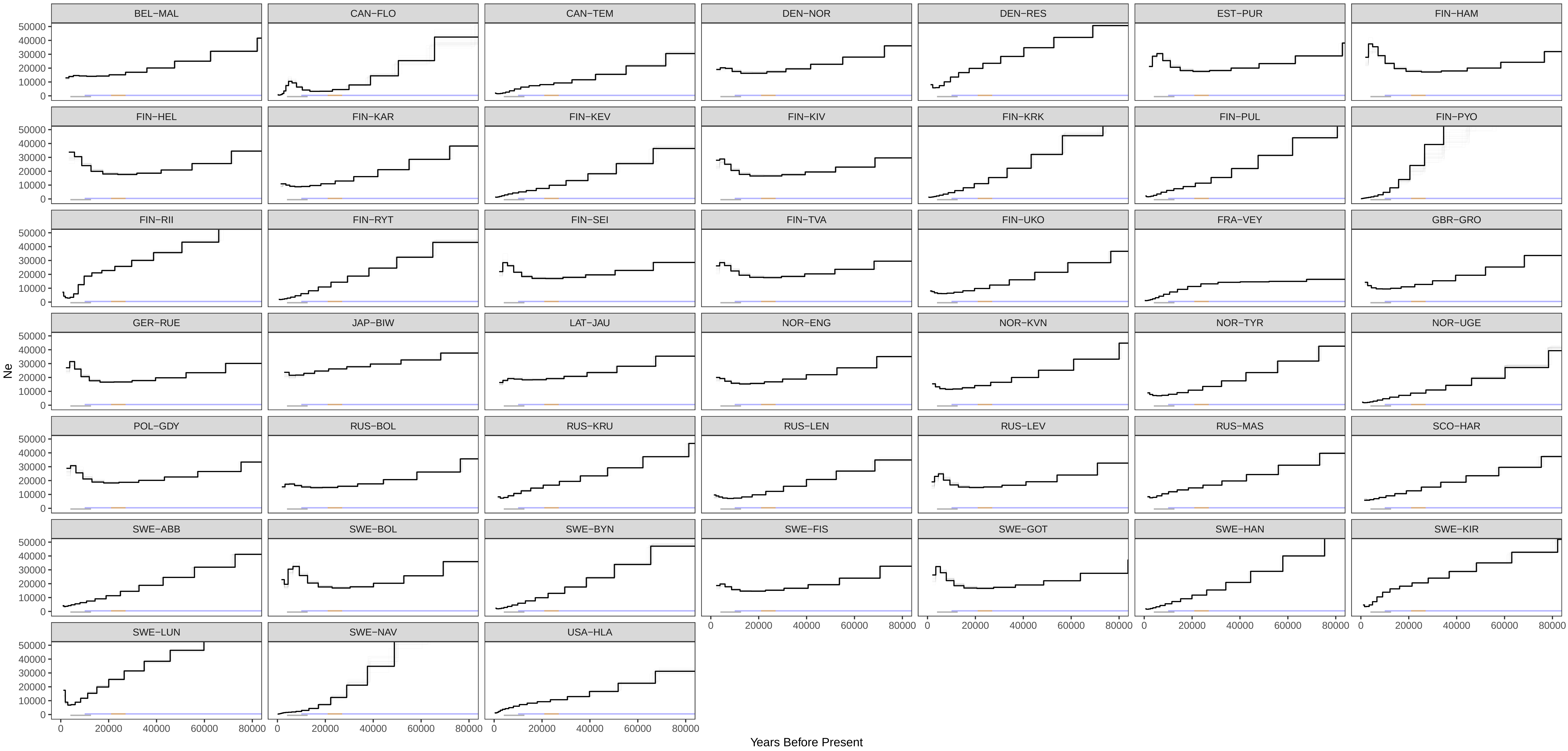

### FigS7.pdf

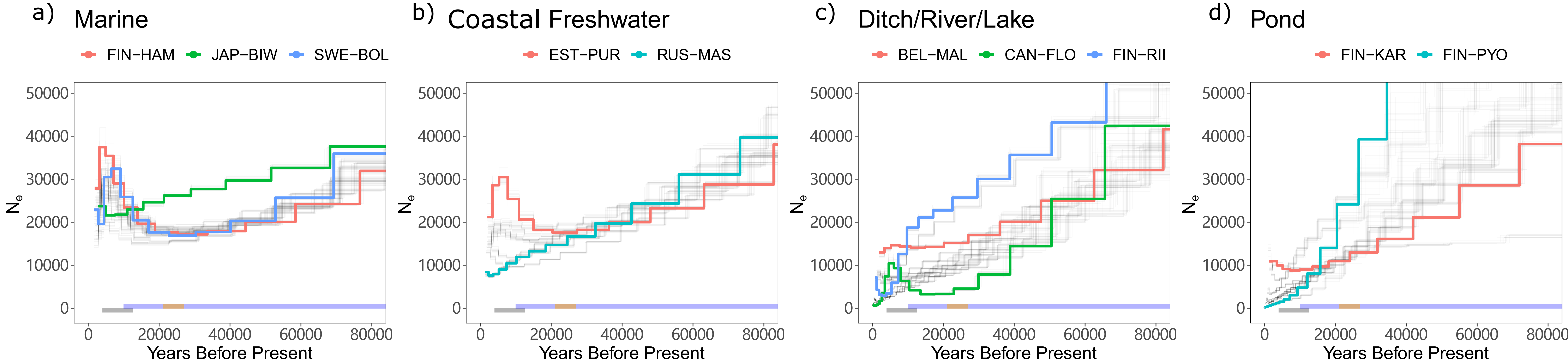

### FigS8.pdf

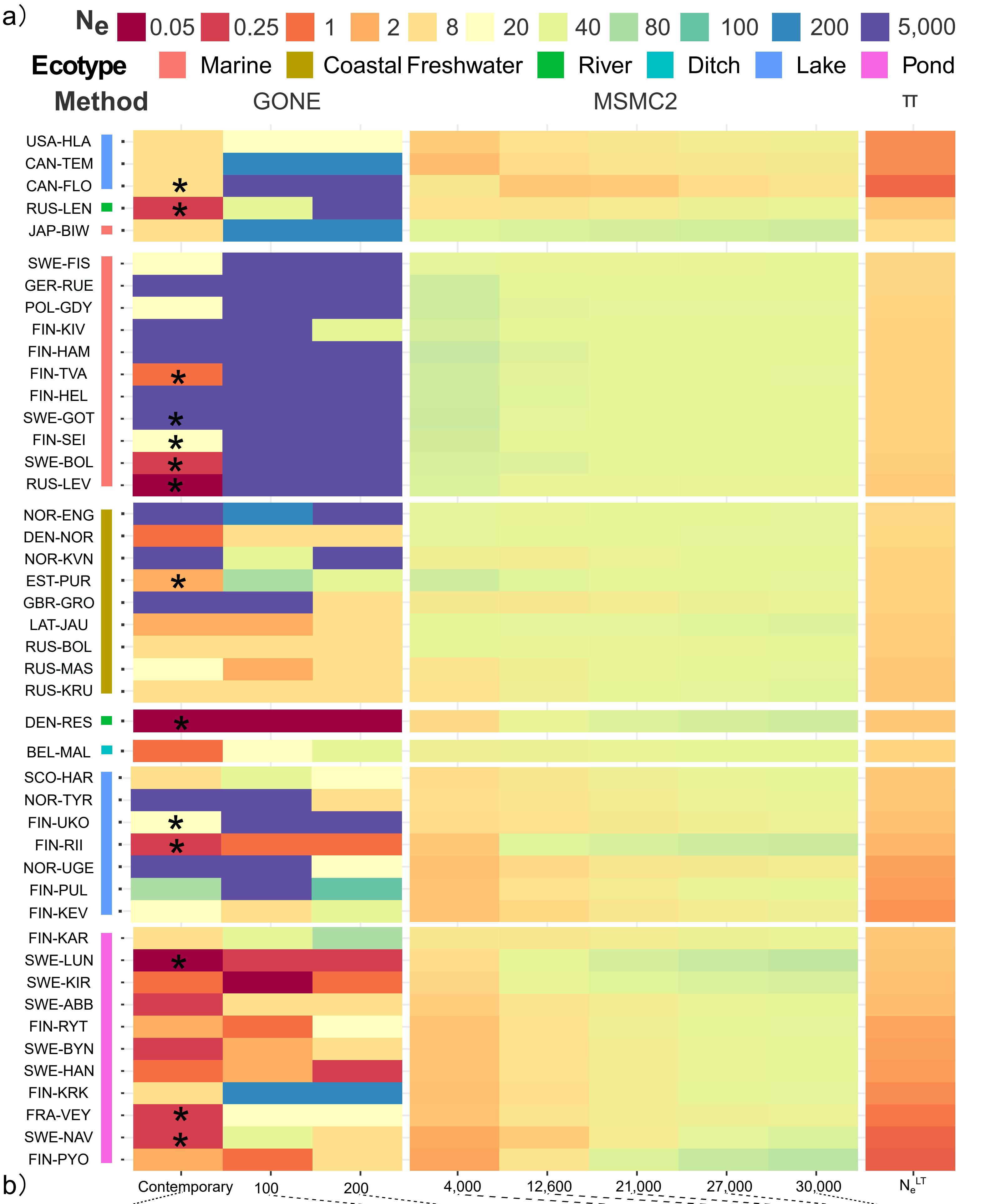

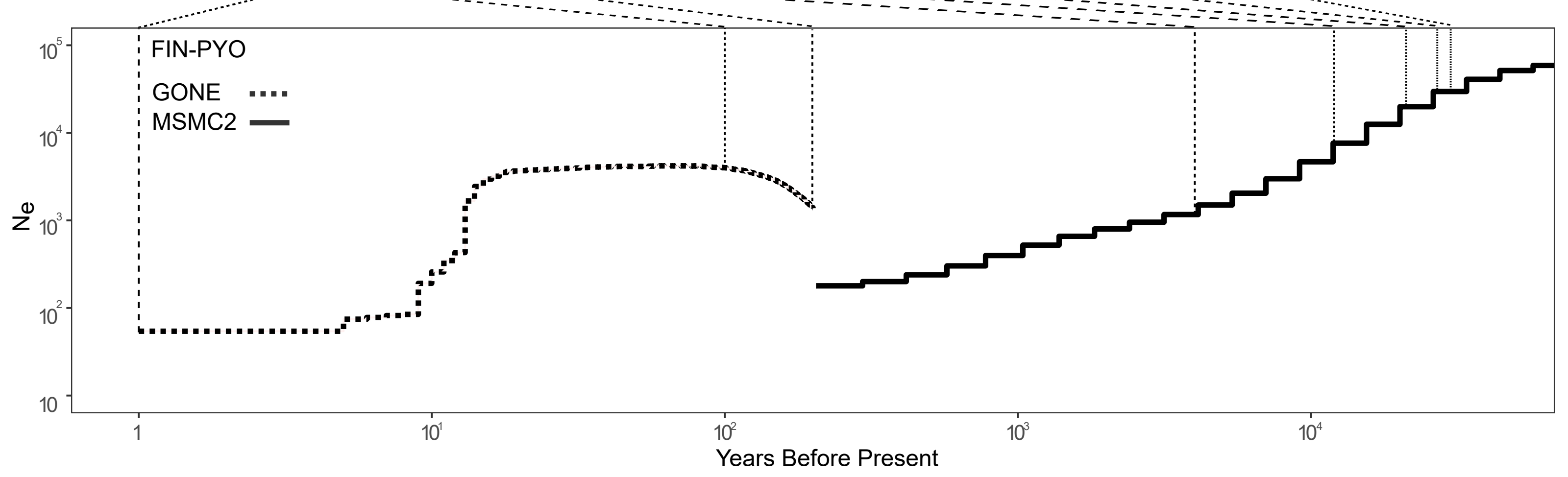
